# Cell-of-Origin Analysis of Metastatic Gastric Cancer Uncovers the Origin of Inherent Intratumor Heterogeneity and a Fundamental Prognostic Signature

**DOI:** 10.1101/725390

**Authors:** Ruiping Wang, Shumei Song, Kazuto Harada, Guangchun Han, Melissa Pool Pizzi, Meina Zhao, Ghia Tatlonghari, Shaojun Zhang, Yuanxin Wang, Shuangtao Zhao, Brian D. Badgwell, Mariela Blum Murphy, Namita Shanbhag, Jeannelyn S. Estrella, Sinchita Roy-Chowdhuri, Ahmed Adel Fouad Abdelhakeem, Guang Peng, George A. Calin, Samir Hanash, Alexander J. Lazar, Andrew Futreal, Jaffer A. Ajani, Linghua Wang

## Abstract

Intra-tumoral heterogeneity (ITH) is the fundamental property of cancer, however, the origin of ITH remains poorly understood. Here we performed single-cell RNA sequencing of peritoneal carcinomatosis (PC) from 20 patients with advanced gastric adenocarcinoma (GAC), constructed a transcriptome map of 45,048 PC cells, determined the cell-of-origin of each tumor cell, and incisively explored ITH of PC tumor cells at single-cell resolution. The links between cell-of-origin and ITH was illustrated at transcriptomic, genotypic, molecular, and phenotypic levels. This study characterized the origins of PC tumor cells that populate and thrive in the peritoneal cavity, uncovered the diversity in tumor cell-of-origins and defined it as a key determinant of ITH. Furthermore, cell-of-origin-based analysis classified PC into two cellular subtypes that were prognostic independent of clinical variables, and a 12-gene prognostic signature was then derived and validated in multiple large-scale GAC cohorts. The prognostic signature appears fundamental to GAC carcinogenesis/progression and could be practical for patient stratification.

---

Gastric adenocarcinoma (GAC) remains a common and lethal disease with a poor prognosis^1^. Often diagnosed at an advanced stage, GAC is frequently resistant to therapy^2^. A common site of metastases is the peritoneal cavity (peritoneal carcinomatosis; PC) and there is an unmet need for improved therapeutic options in advanced GAC patients^3,4^. Patients with PC are highly symptomatic and can have an overall survival of <6 months. Only a small fraction of patients benefits, often only transiently, from immune checkpoint inhibitors^5,6^ or HER2-directed therapy^7^. Molecular understanding of advanced GAC is limited. Four genotypes defined by The Cancer Genome Atlas (TCGA) were based on analysis of primary GACs^8^. The two clinically favorable subtypes, Epstein-Barr virus-induced and microsatellite instable, are rare in advanced GAC cohorts^9^. In the clinic, empiricism prevails as patients are not routinely stratified and rational therapeutic selection is exceedingly limited.

It is well recognized that GAC is endowed with extensive inter- and intra-tumoral heterogeneity (ITH)^8,9^. ITH is fundamental for GAC survival as it confers therapy resistance and is a major obstacle to improving patient outcome. However, the origins of ITH are poorly understood. Deeper understanding of the cellular/molecular basis of ITH could influence how GACs are treated. Single-cell transcriptome sequencing (scRNA-seq) has emerged as a robust and unbiased tool to assess cellular and transcriptomic ITH^10^.

In this study, we incisively explored ITH of PC tumor cells at the single-cell resolution to obtain an improved understanding of the origins of tumor cells that populate and thrive in the peritoneal cavity. We constructed a transcriptome map of 45,048 PC cells, identified and characterized the cell-of-origins of PC tumor cell. This study, uncovers the diversity in tumor cell-of-origins and defines it as a key determinant of ITH in GAC. The links between cell-of-origin and ITH was illustrated at the transcriptomic, genotypic, cell-cycle state, molecular pathways, and phenotypic levels. Finally, the cell-of-origin-based analysis of PC tumor cells led to a 12-gene fundamental signature, which although derived from PC cells, retained its prognostic significance when applied to several independent localized and advanced large-scale GAC cohorts. These results provide an avenue for patient stratification and novel target discovery for future therapeutic exploitation.

## A single-cell transcriptome map of PC

scRNA-seq was performed on freshly isolated ascites cells from 20 GAC patients (**Fig. 1a, Table S1**). Following quality filtering, we acquired high-quality data for 45,048 cells. A multistep approach was applied to identify PC malignant cells and define immune cell types (see **Methods**). We captured 6 main cell types: tumor cells, fibroblasts, and 4 immune cell types, each defined by unique signature genes (**Extended Data Fig. 1**). The immune cells from different patients clustered by cell type, whereas PC malignant cells clustered distinctly by patient. But it was evident that tumor cells from the short-term survivors clustered closely in t-SNE plot (t-distributed stochastic neighbor embedding, **Extended Data Fig. 2**). In this study, we have focused on PC tumor cells (n=31,131). Five patients with too few tumor cells (<50) were excluded from subsequent analysis. To profile the transcriptomic landscape of PC tumor cells, unsupervised cell clustering was carried out, which uncovered 14 unique cell clusters, with differentially expressed genes (DEGs) specifically marking each cell cluster (**Fig. 1b, Extended Data Fig. 3**). It is worth mentioning that tumor cells from patient IP-070 formed two separated clusters (2 and 12). These results indicated a high degree of inter- and intra-tumoral heterogeneities in PC malignant cells.

**Figure 1.**
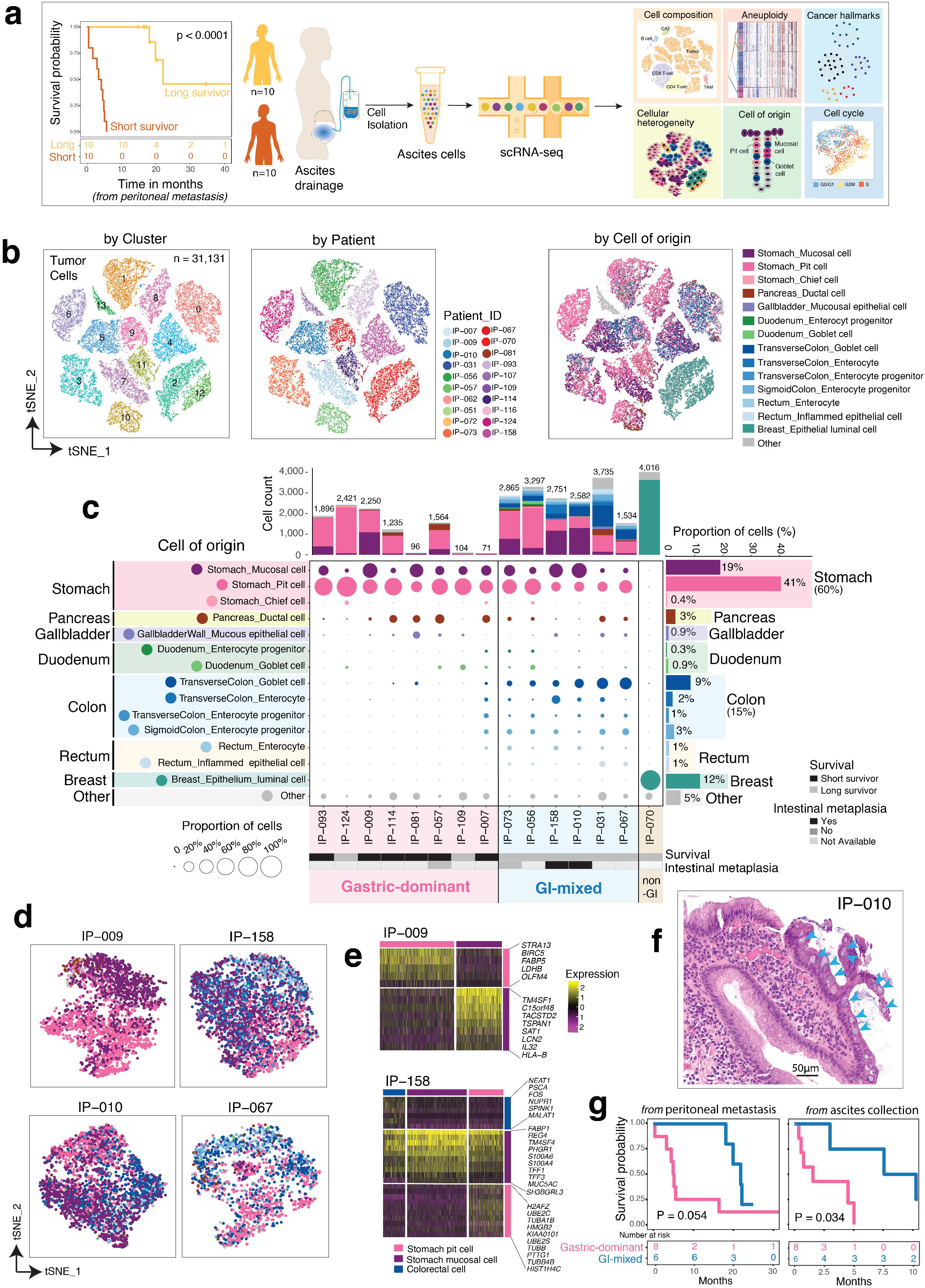
Cell of origin-based classification of gastric peritoneal metastases showed strong correlation with patient survival. This study included 10 short-term survivors and 10 long-term survivors. **a**, (left) The Kaplan-Meier curve demonstrates a dramatic difference in the survival time between two groups of GAC patients; the panels in the middle and right shows a schema of sample collection and scRNA-seq data analysis, respectively. **b**, The tSNE overview of the 31,131 tumor cells (14 cell clusters) that were selected for subsequent analyses in this study. Each dot in the tSNE plot indicates a single cell. Cells are color coded for the tSNE cluster number (left), the corresponding patient origin (middle), and the cell of origin (right). **c**, The cell-of-origin landscape at single-cell resolution showing the origins of PC tumor cells from 15 GAC patients (5 samples were excluded due to less than 50 QC-passed tumor cells with defined cell of origins, Table S2). The middle panel shows the origin (row) of tumor cells by patient (column). The size of the circle represents the proportion of tumor cells (of the total QC-passed tumor cells for each individual sample) for each specific cell origin. The circles are color coded by the cell of origin. The annotation track on the left shows a brief description of each defined cell origin. The histogram on the top shows the number of cells accumulated on 14 listed cell origins (plus Other-other unclassified or rare cell types) in each individual sample (patient). The histogram on the right shows the proportion of tumor cells (of the total QC-passed tumor cells for this cohort) for each specific tumor cell origin. The bottom annotation tracks show (from top to bottom): the corresponding patient ID, classification based on patient survival, the presence of intestinal metaplasia in the corresponding primary tumor, and classification based on the composition of cell compartments. **d**, The representative tSNE plot of tumor cells (colored by their cell of origin) for each individual case, and **e**, The scaled expression values of discriminative genes for each defined tumor cell origin for two representative cases IP-009 and IP-158. **f**, A representative histology image for IP-010 demonstrating well-formed goblet cells in gastric mucosa (blue arrow heads). **g**, The correlation of cell or origin-based classification with patient survival. The survival time was calculated from the diagnosis of peritoneal metastasis (left) and the time of ascites collection (right), respectively.

## The cell-of-origin of malignant cells within the peritoneal cavity

To map each individual PC tumor cell and to determine its cell of origin (genotype/phenotype), we performed cell-of-origin analysis by mapping scRNA-seq data to Human Cell Landscape (http://bis.zju.edu.cn/HCL/), a scRNA-seq database that comprises >630k cells covering 1,393 cell types/states from 44 human organs/tissues (see **Methods**). Our analysis revealed a high degree of cellular heterogeneity in PC (the diversity of origins of tumor cells that comprised the tumor). Intriguingly, although all cases in this study were clinically diagnosed as PC from GAC, our transcriptome-based cell-of-origin analysis revealed 14 defined cell types originated from 7 organs (**Fig. 1c, Table S2**). Only 60% of mapped PC tumor cells transcriptomically resembled cells of stomach origin, including pit cells (41%), mucosal cells (19%), and chief cells (0.4%). However, the expression features of a subset of PC tumor cells (23%) closely resembled cells of other GI (gastrointestinal) organs, including colon (15%), pancreas (3%), rectum (2%), duodenum (1%), and gallbladder (1%). For case IP-070, our analysis suggested that no PC tumor cells were of GI origin, instead, the cells transcriptomically resembled breast luminal epithelial cells (**Extended Data Fig. 4**). After a comprehensive review of the patient’s clinical record, we noted that this case was misdiagnosed and treated as GAC at an outside hospital but it was breast cancer that metastasized to the stomach resulting in PC subsequently. This vignette reflected the accuracy of our cell-of-origin analysis.

## The diversity in tumor cell origins is a key determinant of PC transcriptomic ITH with prognostic value

To further study ITH and examine its relationship with tumor cell origins, we performed unsupervised clustering of PC tumor cells separately for each individual case based on transcriptomic profiles and then projected the cell-of-origin annotation on generated tSNE plots. The representative results are shown in **Fig. 1d**. We observed a separation (different transcriptomic profiles) of the cells showing a gastric lineage from the cells demonstrating a colonic lineage (IP-067, IP-158, IP-010) in tSNE plots. Notably, the stomach pit cells also clustered distinctly from stomach mucosal cells (IP-009). DEGs analysis revealed gene expression signatures specific to each cell population (**Fig. 1e**). Our results demonstrated that PC tumor cells with different cell origins are transcriptomically distinct and suggested that the diversity in tumor cell origins is likely a determinant of ITH. For two cases (IP-158 and IP-010) with mixed stomach/colon cell lineages, we were able to retrieve the histology images of the corresponding GAC primary tumors and confirmed tumors arose in the setting of gastric intestinal metaplasia, characterized by the presence of well-formed goblet cells in gastric mucosa (**Fig. 1f**). This finding is intriguing given the associated analysis showing a mixed cellular population of both gastric and colorectal lineages.

Based on the cellular compositions, we classified PC into two main groups: the Gastric-dominant (dominated by gastric cell lineages) and the GI-mixed (with mixed gastric and colorectal cell lineages) (**Fig. 1c**). We further investigated the correlation of cell-of-origin-based classification with clinical/ histopathological variables, and no significant difference was observed for histopathological features between two groups. Notably, the cell-of-origin based classification of PC tumor cells showed a strong correlation with patient survival (**Fig. 1g**): all 6 cases with a GI-mixed phenotype were long survivors, whereas 6/8 cases with a Gastric-dominant phenotype were short survivors (Fisher’s Exact test, P = 0.0097, **Extended Data Fig. 5**). Currently, a validated and practical molecular signature for PC is lacking. These results suggested that, the cell-of-origin features of PC tumor cells could prognosticate patient survival.

## Tumor cell proliferative property strongly correlated with tumor cell-of-origin

To study the ITH of tumor cell proliferative property and examine its link to tumor cell-of-origin, we computationally assigned a cell cycle stage to each individual cell based on expression profile of cell-cycle related signature genes^11^ (see **Methods**). Our analysis suggested that 51% of PC tumor cells are cycling, either in G2M or S phase (**Fig. 2a, Table S3**). Interestingly, tumor cell proliferative property strongly correlated with tumor cell origins (**Fig. 2b**). The stomach pit cells were highly proliferative, with vast majority of cells in G2M/S phase, while the stomach mucosal cells and cells of colorectal origins were quiescent. Consistently, some key cell-cycle regulatory genes were differentially expressed across tumor cell populations with different origins and associated with patient survival (**Fig. 2c**).

**Figure 2.**
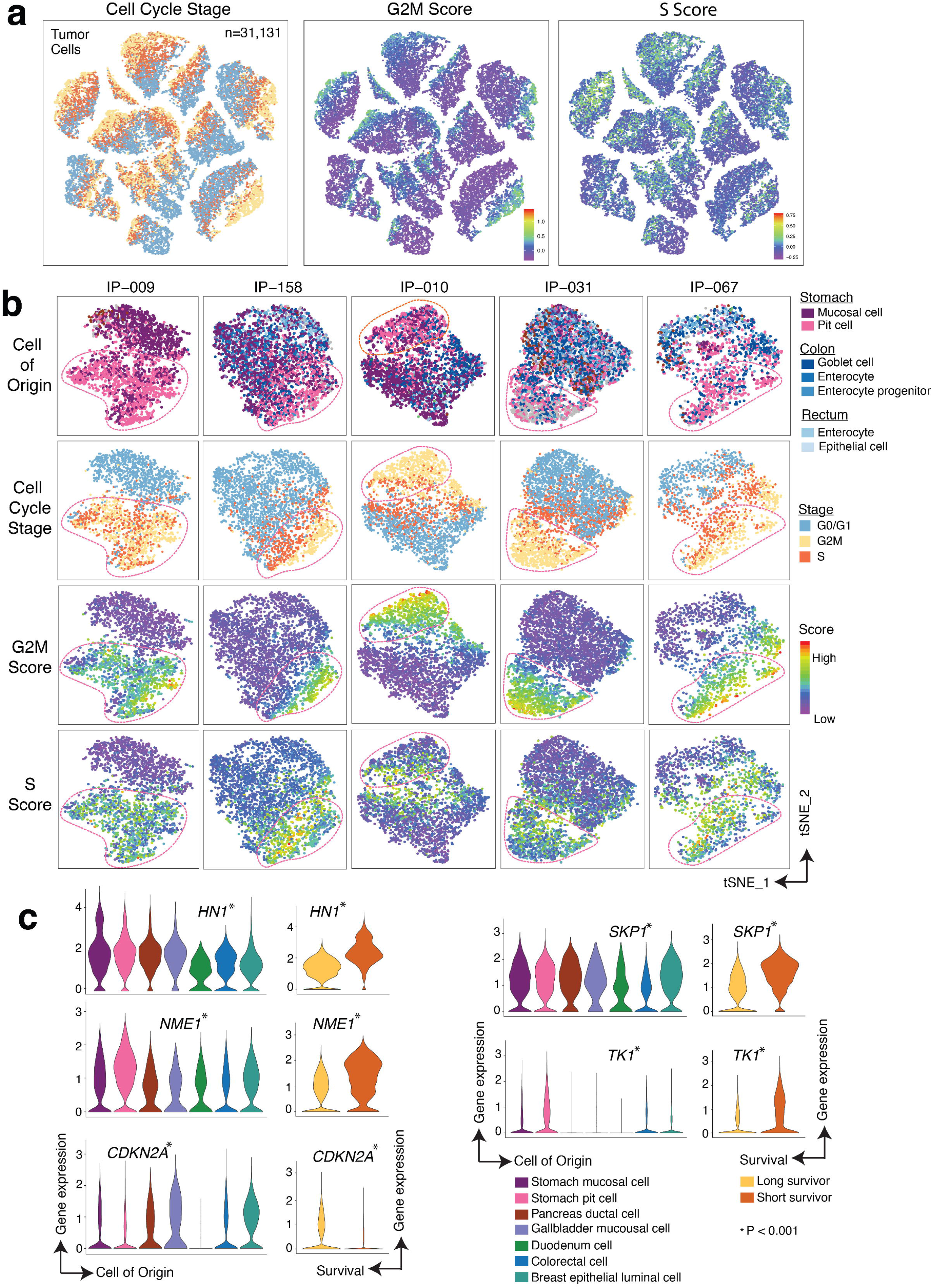
The tumor cell transcriptome heterogeneity and cell proliferative properties closely associated with tumor cell origins. **a,** The tSNE overview of the cell cycle stages (left), the G2M score (middle), and S score (right) for the 31,131 QC-passed PC tumor cells. **b,** The tSNE plots of 5 representative cases displaying unsupervised clustering of tumor cells according to their transcriptome profiles and the correlation of cell proliferative property with tumor cell origins. The cells are colored coded (from top to bottom) by tumor cell-of-origin, cell cycle stage, the quantitative score for G2M phase and S phase, respectively. The red irregular shapes are used to highlight the highly proliferative cell cluster of each individual sample. **c,** The violin plots for representative genes that are differentially expressed between tumor cells with different cell origins (left) and their correlation with patient survival (right).

## The Genotypic ITH of PC tumor cells links to cell-of-origin

We next investigated the genotypic ITH of PC tumor cells and examined its association with tumor cell origins. Single-cell copy number variations (CNVs) were inferred from scRNA-seq data^10,12,13^ (see **Methods**). The inferred CNVs showed considerable patient-to-patient and cell-to-cell variations (**Fig. 3**), indicating a significant genotypic heterogeneity among PC tumor cells. For each individual patient, we further investigated copy number subclonal structures of PC tumor cells using unsupervised hierarchical clustering. Intriguingly, the pattern of CNV subclonal structures aligned well with tumor cell origins (**Fig. 3a**). For example, for case IP-067, PC tumor cells clustered into 3 major subpopulations with distinct CNV profiles: the largest subpopulation was mainly comprised of cells of colon lineage and distinguished by number of CNVs from the smallest subpopulation that was purely comprised of cells of stomach lineage, a subpopulation in medium size was a mixture of cells from both lineages and the cells shared similar CNV profiles. Similarly, 3 populations were identified in IP-009 that was gastric-dominant (**Fig. 3a**, bottom), and the smallest subpopulation (comprised of stomach pit cells) showed additional CNVs that were not present in the subpopulation that was mainly comprised of stomach mucosal cells.

**Figure 3.**
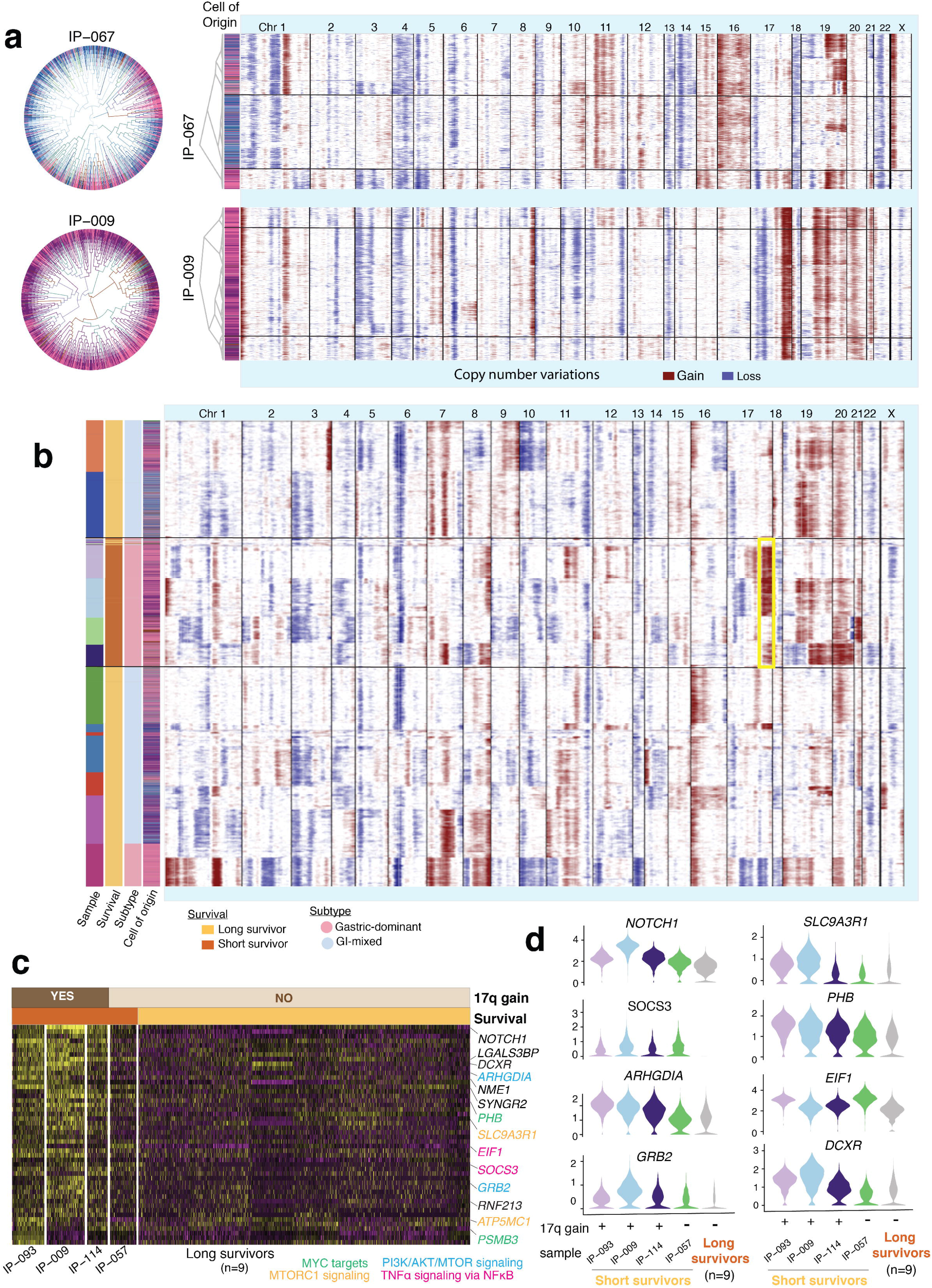
DNA copy number variations (CNVs) and genotypic heterogeneity associated with tumor cell origin and patient survival. **a,** An overview of the genome-wide CNVs for two representative cases (IP-067, GI-mixed; IP-009, Gastric-dominant) and correlation of the genotypic heterogeneity with the origins of tumor cells. The circus plots on the left demonstrated unsupervised clustering of tumor cells with different origin (color coded) for each individual case based on their inferred CNVs. The unsupervised hierarchical clustering on the right displays a detailed map of the CNVs across 22 chromosomes (labelled on the top) for each individual cell (row), with copy number gains in red and losses in blue. The dendrogram on the left indicates the clustering structure and the annotation track next to it shows the defined cell of origin (color coded as in Fig. 1c). **b,** The landscape of inferred CNVs for all 31,131 tumor cells. The annotation tracks on the left indicates the corresponding patient ID, survival status, classification based on cell of origin, and tumor cell origins, respectively. The chromosome numbers were labelled on the top. The yellow rectangle highlights the 17q copy number gain that was observed exclusively in cells from the short-term survivors. **c,** The heatmap displays scaled expression values of genes upregulated in 3 short survivors (sample IDs labelled at the bottom) with evident 17q gain (annotated on the top track), 1 short-term survivor and 9 long-term survivors without detectable 17q changes. Biologically important genes were listed on the right, color coded by their related signaling pathways. **d,** The representative violin plots of 8 genes selected from the Panel **c**.

In addition, we analyzed CNVs from all cases together and discovered 17q amplification as a unique event that was highly abundant in tumors cells with stomach origin and only present in cells from short survivors (**Fig. 3b**). By integrating genotypic and transcriptomic profiles, we identified a list of upregulated genes on 17q in tumor cells with evident 17q amplification (compared to the rest of cells without amplified 17q) (**Fig. 3c, Table S4**). Some of these upregulated genes involved in key signaling pathways (PI3K/AKT/mTOR, mTORC1, MYC), are potential therapeutic targets (*NOTCH1, GRB2, PSMB3*) with a number of compounds being screened as active^14^ (**Table S5**), and associated with patient survival (**Fig. 3d**). Our results demonstrated that the genotypic ITH in PC tumor cells associated with tumor cell origin and patient survival.

## Single-cell molecular signaling heterogeneity correlated with tumor cell-of-origin

To examine the molecular consequences of transcriptomic and genotypic alterations described above and to better understand the biological programs associated with cell-of-origin and patient survival, we performed integrative analysis of >900 molecular signaling pathways. Among them, 80 pathways were differentially expressed across tumor cell origins (**Fig. 4a**), and of these, 37 were also strongly associated with patient survival (**Fig. 4b, Extended Data Fig. 6**). These pathways were categorized into 5 major classes based on their biological functions: oncogenic signaling, cell cycle, DNA repair, metabolism, and immune signaling (**Fig. 4c**). Pathway interactions analysis revealed that these biological processes are functionally connected.

**Figure 4.**
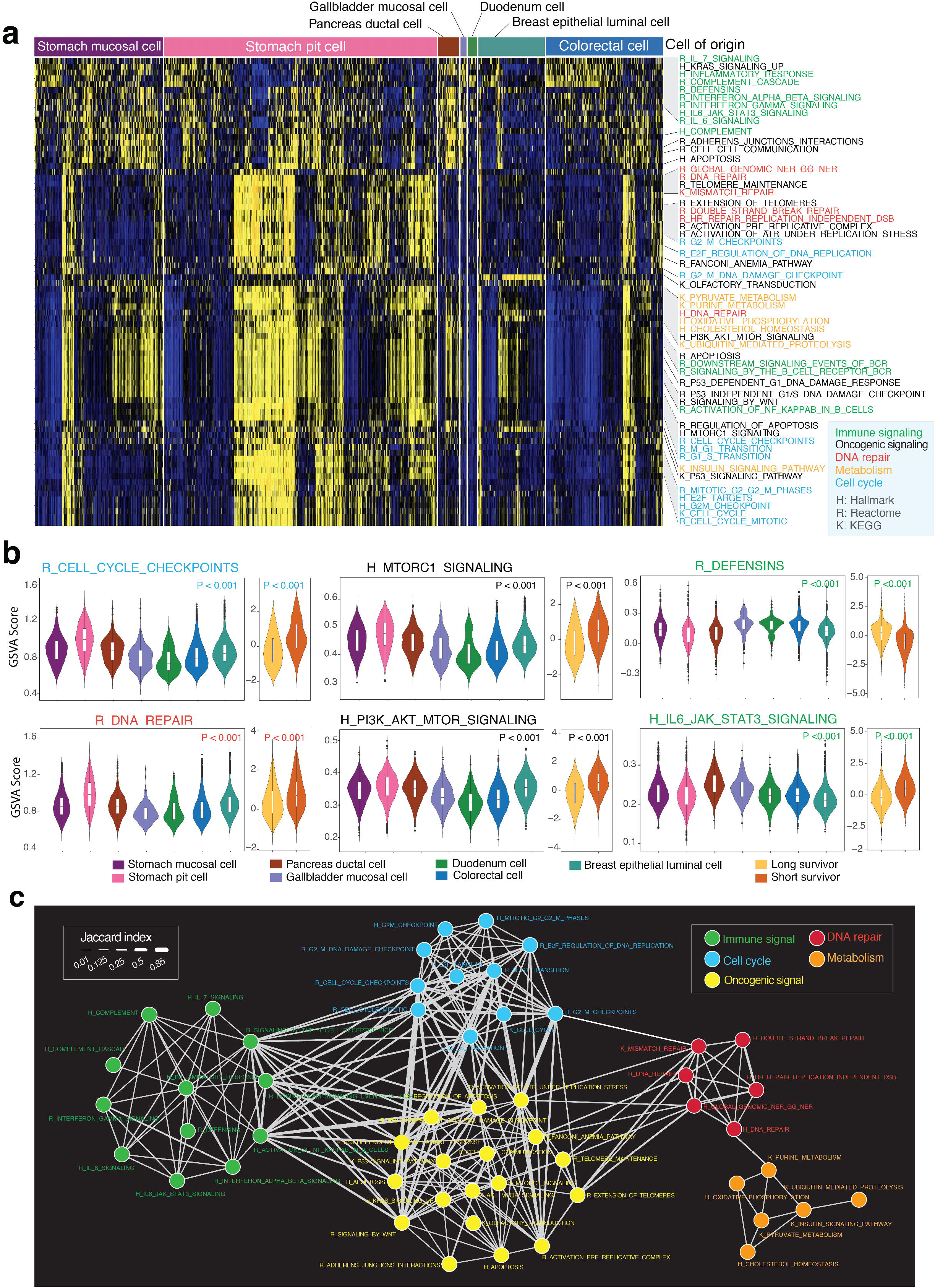
Molecular pathway based dissection of the transcriptomic heterogeneity and correlation with cell-of-origin and patient survival. **a,** The transcriptomic heterogeneity of annotated gene sets including cancer hallmark gene sets (n=50), and other curated gene sets from KEGG (n=186) and Reactome (n=674) pathway databases. Each column represents a single cell. Only the pathways (row) that differentially expressed across different tumor cell origins are shown. The tumor cell origin was annotated at the top track and the pathway names are labelled on the right, color coded by their biological functions. **b,** The representative violin plots of 6 pathways selected from Panel **a** and Extended Data Fig. 6 that showed significant correlation with patient survival. **c,** The interaction networks of differentially expressed pathways displayed in the Panel **a**.

Pathways that were significantly enriched in tumor cells with gastric lineage and associated with shorter survival included cell cycle, DNA repair, PI3K/AKT/mTOR, mTORC1, Wnt, NFκB, and metabolic reprogramming, which are predominantly oncogenically encoded. In contrast, pathways that were enriched in tumor cells with colon lineage and associated with longer survival included defensins, IL-7 signaling, complement cascade, IL6/JAK/STAT3 signaling, and interferon alpha/gamma, which are exclusively immune related (**Figs. 4a-b**). These results indicated that different biological processes might have been activated in tumor cells with different origins and contributed to their distinct molecular consequences and patient survival.

## Generation and validation of a cell-of-origin based 12-gene prognostic signature

Based on cell-of-origin analysis, a 12-gene signature was derived (**Figs. 5a-b, Methods**). We first validated this signature in an independent, advanced GAC cohort (n=45) using bulk RNA-seq data. This signature demonstrated a great power to prognosticate patient survival and consistently, patients with a Gastric-dominant molecular feature in their PC cells survived significantly shorter (7.8 vs. 24.5 month) than those with a GI-mixed feature (**Fig. 5c**). Multivariable Cox regression analysis showed that this signature is a strong prognosticator of short survival and it outperformed all clinical variables and was independent of clinical/histopathological features (**Extended Data Fig. 7**).

**Figure 5.**
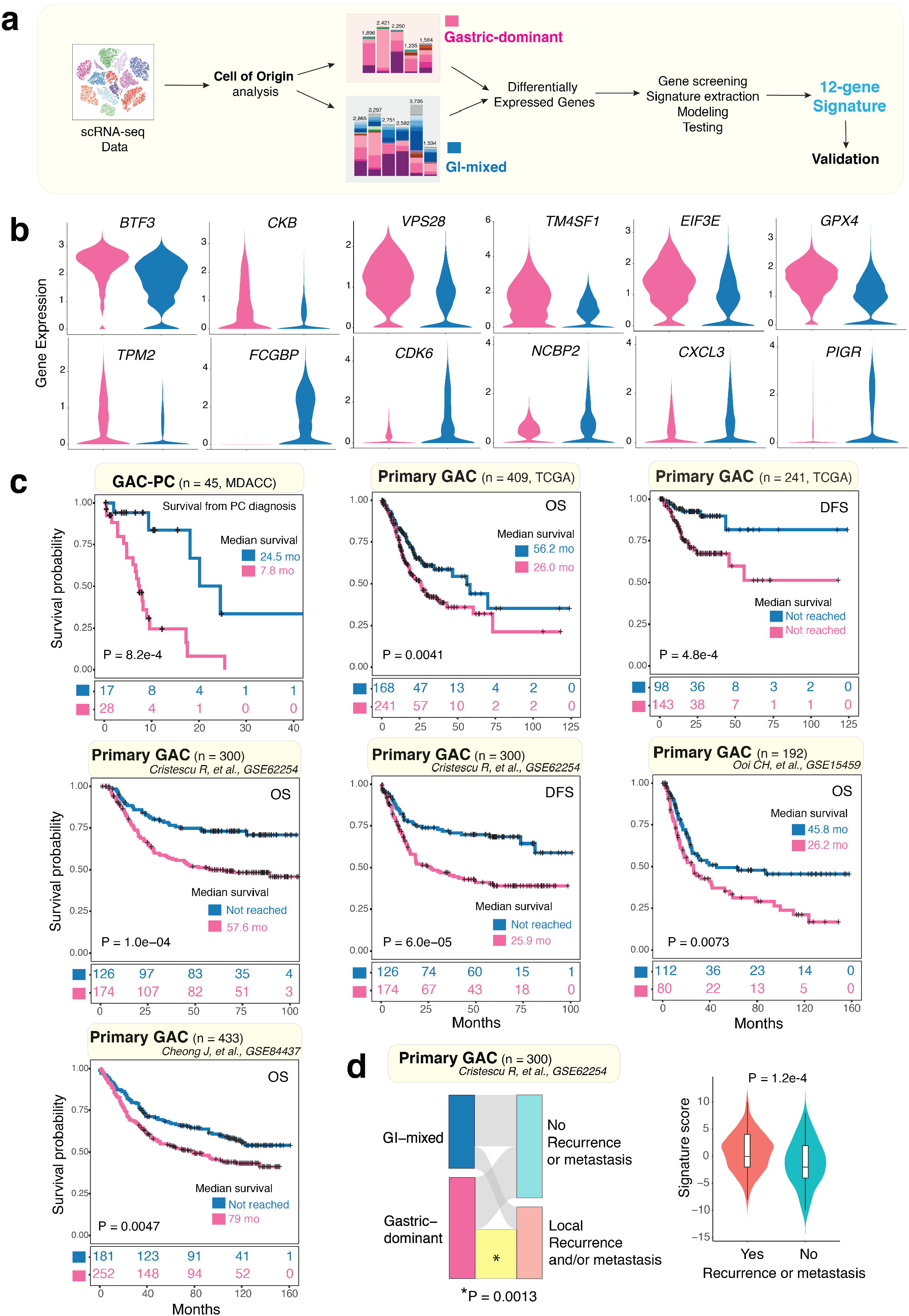
Identification and validation of the 12-gene prognostic signature. **a,** A schema that describes the bioinformatics flow for generation of the 12-gene signature. **b,** The 12 genes used to generate this signature and its differential expression between the gastric-dominant and GI-mixed groups. **c,** The Kaplan–Meier curves demonstrating the predictive power of this signature across 5 validation cohorts including a second independent cohort of GAC-PC patients (n=45) from MD Anderson Cancer Center (MDACC), the TCGA primary GAC cohort, and 3 other large-scale primary GAC cohorts. The source of the dataset, the size of each cohort, log-rank P-value, and the median survival time (in months) were labeled on each Kaplan–Meier plot. OS, overall survival; DFS, disease-free survival. **d,** The alluvial plots (left) shows the relationship between cell-of-origin defined subtypes (left strip) and the presence of local recurrence and/or distal metastasis (right strip). The yellow band highlights the significant enrichment of local recurrence and/or distal metastasis events in tumors with the gastric-dominant subtype. The violin plot (right) shows a significant difference in the mean signature score in tumors with/without local recurrence and/or distal metastasis.

We next evaluated its prognostic significance in 5 other large-scale localized GAC cohorts^13,15–17^, totaling 1,425 patients. Notably, although this signature was derived from an advanced GAC cohort, it retained its prognostic prowess in these validation cohorts of localized GACs and demonstrated a robust power in prognosticating survival (**Fig. 5c**). Intriguingly, this signature is independent of other molecular and clinical subtypes (**Extended Data Fig. 8**) and it correlated strongly with the risk of local recurrence/distal metastases among the TCGA^8^ and Cristescu cohorts^17^, where expression and outcome data are both available (**Fig. 5d, Extended Data Fig. 9**). These results further highlighted the value of this prognostic signature and its robustness in prognosticating patient survival.

## Discussion

The progress against GAC has lagged behind other GI tumor types. Therapy resistance and the lack of rational therapeutic targets against GAC represent major obstacles in improving survival of advanced GAC patients^18^. It is widely appreciated that ITH is a fundamental property of cancer contributing to therapeutic failure, development of distant metastases,^19^ and hindrance to biomarker/target discoveries^20^. Recent studies of localized and advanced GACs identified multiple molecular subtypes and revealed a high degree of ITH that are associated with poor clinical outcomes^9,21,22^. Therefore, deeper dissection of ITH is critical for understanding the underlying mechanisms driving poor prognosis of GAC and for overcoming therapeutic resistance. In this study, we dissected, at unprecedented resolution, the cellular and transcriptomic ITH of PC tumor cells using the cutting-edge scRNA-seq technology, in combination with integrative computational analyses.

A key finding of this study is that diversity of cell-of-origin appears to mirror and may even dictate inherent ITH of PC tumor cells at multiple molecular levels. The origin of ITH has been the subject of discussion, with multiple models being proposed^23,24^. Peritoneal cavity is a unique microenvironment where tumor cells can be in suspension in the peritoneal fluid as opposed to localized solid tumor tissues, the ascites cells we have sequenced may be a better representation of ITH. We discovered several transcriptomically distinct tumor cell populations that could be distinguished by cell lineage characteristics. We noted that >40% of cases in our discovery cohort had a large number of tumor cells with genotype/phenotype mapping to non-stomach GI lineages. We documented that ITH defined by cell-of-origin is perpetuated at transcriptomic, genotypic, cell-cycle state, molecular signaling, and phenotypic levels and strongly associated with survival. We showed that tumor cell transcriptomic profiles and proliferative property significantly differed across cells with different origins, so did the molecular signaling, suggesting that treatment strategies could potentially be tailored to these molecular features. It would appear that varied biological programs (e.g. genomic/epigenomic) might have been engaged early in tumor cells resulting in different genotypes/phenotypes and subsequently contributed to distinct molecular ITH and patient prognosis. In addition, we discovered that 17q amplification was highly abundant in PC cells of gastric lineage. 17q is a region that harbors multiple potential therapeutic targets and interestingly, all patients with 17q amplification had a short survival. Our discovery of the direct link between tumor cell-of-origin and ITH at the single-cell resolution could be generalized to other cancer types and broaden our understanding of cancer in general.

Most intriguingly, the cell-of-origin-based analysis classified PC tumor cells into two cellular subtypes that were prognostic independent of histopathological features. Further analyses led us to discover a 12-gene signature that appears to be fundamental to GAC carcinogenesis/propagation as it was not only highly prognostic in GAC metastatic validation cohort but perform just as robustly in several large-scale localized GAC cohorts. Currently, there is no such signature in clinical use and this signature has a high potential to stratify patients for more effective therapies as this becomes available.

## METHODS

### Patient cohort, clinical characteristics, and sample collection

A total of 20 GAC patients with malignant ascites (peritoneal carcinomatosis, PC) was included in this study. The detailed clinical and histopathological characteristics are described in the **Supplementary Table S1**. GACs were staged according to the American Joint Committee on Cancer Staging Manual (8th edition)^25,26^. PC was confirmed by cytologic examination. This cohort included 10 long-term survivors and 10 short-term survivors. The long-term survivors were patients who survived more than 1 year after the diagnosis of PC and the short-term survivors were patients who passed away within 6 months after the diagnosis of PC. Based on the Lauren’s classification of the primary GAC, all tumors were of diffuse type. Sixteen out of twenty patients had Signet-ring cell carcinoma. Her 2 positivity was performed but no Her2 positivity was detected in these patients. PC specimens were collected at The University of Texas MD Anderson Cancer Center (Houston, USA) under an Institutional Review Board (IRB) approved protocol after obtaining written informed consent from each participant. Patients with diagnosed GAC-PC with ascites were approached when they required a therapeutic paracentesis. No other selection criteria were applied. This project was approved by the IRB and is in accordance with the policy advanced by the Helsinki Declaration of 1964 and later versions. PC specimens were spun down for 20 minutes at 2,000g and pelleted cells (PC cells) were isolated, property store at −80 °C and used for scRNA-seq. To minimize batch effects, the samples were processed together using the same protocol by the same research assistant.

### scRNA-seq library preparation and sequencing

Chromium™ Single cell sequencing technology from 10X Genomics was used to perform single cell separation, cDNA amplification, and library construction following the manufacturer’s guidelines. Briefly, the cellular suspensions were loaded on a 10x Chromium Single Cell Controller to generate single-cell Gel Bead-in-Emulsions (GEMs). The scRNA-Seq libraries were constructed using the Chromium Single Cell 3’ Library & Gel Bead Kit v2 (PN-120237, 10x Genomics). The HS dsDNA Qubit kit was used to determine concentration of both the cDNA and libraries. The HS DNA Bioanalyzer was used for quality track purpose and size determination for cDNA and lower concentrated libraries. Sample libraries were normalized to 7.5 nM and equal volumes added of each library for pooling. The concentration of the library pool was determined using Library Quantification qPCR kit (KAPA Biosystems) prior to sequencing. The barcoded library at the concentration of 275 pM was sequenced on the NovaSeq6000 (Illumina, San Diego, CA), S2 flow cell (100 cycle kit) using a 26 × 91 run format with 8 bp index (read 1). To minimize batch effects, all sequencing was processed together as a single batch. The libraries were constructed using the same version of reagent kits following the same protocols and the libraries were sequenced on the same flow cell and analyzed together.

### scRNA-seq data processing and analysis

#### Raw sequencing data processing, QC, data filtering, and normalization

The raw single cell RNA sequencing data were pre-processed (demultiplex cellular barcodes, read alignment, and generation of gene count matrix) using Cell Ranger Single Cell Software Suite provided by 10x Genomics. Detailed QC metrics were generated and evaluated. Genes detected in <3 cells and cells where < 200 genes had nonzero counts were filtered out and excluded from subsequent analysis. Low quality cells where >15% of the read counts derived from the mitochondrial genome were also discarded. After applying these QC criteria, 45,048 single cells and 23,057 genes in total remained and were included in subsequent downstream analysis. Possible batch effects were evaluated using principal component analysis (PCA). In this study, all sequencing libraries were constructed using the same version of reagent kits following the same protocols and the libraries were sequenced on the same illumine platform. Therefore, no significant batch effects were observed. Library size normalization was performed in Seurat^27^ on the filtered gene-cell matrix to obtain the normalized UMI count as previously described^28^.

#### Unsupervised cell clustering and dimensionality reduction

Seurat^27^ was applied to the normalized gene-cell matrix to identify highly variable genes for unsupervised cell clustering. To identify highly variable genes, the *MeanVarPlot* method in the Seurat^27^ package was used to establish the mean–variance relationship of the normalized counts of each gene across cells. We then chose genes whose log-mean was between 0.0125 and 3 and whose dispersion was above 0.5, resulting in 3,018 highly variable genes. The elbow plot was generated with the *PCElbowPlot* function of Seurat^27^ and based on which, the number of significant principal components (PCs) were determined. Different resolution parameters for unsupervised clustering were then examined in order to determine the optimal number of clusters. For this study, the first 10 PCs and the highly variable genes identified by Seurat^27^ were used for unsupervised clustering with a resolution set to 0.6, yielding a total of 20 cell clusters. The t-distributed stochastic neighbor embedding (t-SNE) method was used for dimensionality reduction and 2-D visualization of the single cell clusters.

#### Determination of major cell types and cell states

To define the major cell type of each single cell that mapped to the tSNE plot, feature plots were firstly generated for a suggested set of canonical immune and stromal cell marker genes^29,30^. Enrichment of these markers in certain clusters was considered a strong indication of the clusters representing the corresponding cell types. In addition, differentially expressed genes (DEGs) were identified for each cell cluster using the *FindAllMarkers* analysis in the Seurat^27^ package, followed by a manual review process. The two approaches are combined to infer major cell types for each cell cluster according to the enrichment of marker genes and top-ranked differentially expressed genes in each cell cluster, as previously described^30^.

#### Infer large copy number variations, distinguish tumor cells

InferCNV was applied to infer the large-scale copy number variation (CNVs) from scRNA-seq data (inferCNV of the Trinity CTAT Project; https://github.com/broadinstitute/inferCNV) and the monocytes from this dataset were used as the control for CNVs calling. Initial CNVs were estimated by sorting the analyzed genes by their chromosomal locations and applying a moving average to the relative expression values, with a sliding window of 100 genes within each chromosome, as previously described^10,13^. Malignant cells were distinguished from normal cells based on genomic CNVs, inferred aneuploidy status, cluster distribution of the cells, and marker genes expression.

#### Cell of origin analysis

The cell of origin was assigned by cell type mapping R package scHCL(https://github.com/ggjlab/scHCL) by mapping our transcriptomic data to scHCL (a scRNA-seq database that comprises >630K single cells covering 1,393 cell types/states from 44 human organ and tissue types) and identifying the best match (Spearman’s rank-order correlation) for each cell.

#### Inferring cell cycle stage, hierarchical clustering, differentially expressed genes (DEGs), and pathway enrichment analysis

The cell cycle stage was computationally assigned for each individual cell by the function *CellCycleScoring* that is implemented in Seurat^27^. Cell cycle stage was inferred based on expression profile of the cell cycle related signature genes, as previously described^11^. Hierarchical clustering was performed for each cell type using the Ward’s minimum variance method. Differentially expressed genes (DEGs) were identified for each cluster using the *FindMarkers* function of in Seurat R package^27^ and DEG list was filtered with the following criteria: the gene should expressed in 20% or more cells in the more abundant group; expression fold change >1.5; and FDR q-value <0.05. Heat map was then generated using the *heatmap* function in pheatmap R package for filtered DEGs. For pathway analysis, we applied single-sample GSVA (ssGSVA) to determine the molecular phenotypes of single cells using scRNA-seq expression data. The curated gene sets (including Hallmark, KEGG, REACTOME gene sets, n=910) were downloaded from the Molecular Signature Database (MSigDB, http://software.broadinstitute.org/gsea/msigdb/index.jsp), and pathway scores were calculated for each cell using *gsva* function in GSVA software package ^31^. Pathway enrichment analysis was done with the limma R software package. Significant signaling pathways were identified with a FDR q-value < 0.01.

### Datasets

In addition to the scRNA-seq dataset generated internally for this GAC PC cohort, we included the bulk mRNA-seq data generated on an independent GAC PC cohort from our recent study^9^ to validate the 12-gene prognostic signature. Moreover, we downloaded the bulk mRNA-seq expression data (normalized) generated by The Cancer Genome Atlas (TCGA) on primary stomach adenocarcinoma from NCI Cancer Genomic Data Commons (NCI-GDC: https://gdc.cancer.gov). The mRNA-seq expression data was processed and normalized by the NCI-GDC bioinformatics team using their transcriptome analysis pipeline and we downloaded the normalized expression data. The clinical annotation of TCGA patients were downloaded from a recent PanCanAtlas study^32^. Furthermore, we downloaded 5 large-scale primary GAC datasets (GSE14208, GSE62254, GSE15459, GSE84437) from the Gene Expression Omnibus (GEO) database (GEO, https://www.ncbi.nlm.nih.gov/geo/) to further evaluate the prognostic power of our identified 12-gene signature.

### Generation and validation of the 12-gene prognostic signature

To generate a gene expression signature that is clinically applicable, we performed multiple-step analysis (**Fig. 5a**). First, we compared the gene expression profiles of the cell-of-origin based classification of two patient groups (gastric-dominant vs. GI-mixed) and identified differentially expressed genes (DEGs) between the two groups. Only the DEGs that are highly expressed (normalized UMI count >1) in at least 50% of cells from one of the two groups were taken into subsequent analysis. We next screened each DEG based on their statistical correlation with patient survival and only the DEGs showed a significant (P<0.05) (or a clear trend, P<0.15) correlation with patient survival were selected, followed by model testing of all possible multiple gene combinations. The signature was then extracted and subject to validation with both internally generated and publicly available datasets. To select the optimal classification threshold for tumor classification, use the signature score, we tested different possible signature score values and the threshold value 0 was selected. Tumors with a signature score value >0 were classified as gastric-dominant, and tumors with a signature score value <0 were classified as GI-mixed. Higher signature scores correlate with the gastric-dominant phenotype and with worse prognosis. For the bulk expression datasets, the signature scores were calculated using the normalized gene expression values, taking into consideration of the direction of association (a positive score is assigned for genes associated with the gastric-dominant subtype and a negative score is assigned for genes associated with the GI-mixed subtype). The signature scores were further normalized for subsequent survival analysis.

### Statistical analysis

In addition to the bioinformatics approaches described above for scRNA-seq data analysis, all other statistical analysis was performed using statistical software R v3.5.2. Analysis of differences on a continuous variable (such as gene expression, pathway score) across two groups (a categorical independent variable, such as gastric-dominant vs. GI-mixed) was determined by the nonparametric Mann-Whitney U test. The nonparametric Kruskal-Wallis test was applied to assess the significant difference on a continuous variable by a categorical independent variable with multiple groups (such as the different tumor cell of origin groups). <u>Survival analysis</u>: For survival analysis including overall survival (OS), Progression-free interval (PFS), disease-free survival (DFS), disease-specific survival (DSS), disease-free interval (DFI), and survival time from peritoneal metastasis, we used the log-rank test to calculate p-values, between groups, and the Kaplan-Meier method to plot survival curves. For the TCGA dataset, the clinical annotation and the times calculated for OS, DFS, DSS, DFI were downloaded from the PanCanAtlas study^32^. For other large-scale primary GAC datasets downloaded from GEO, the relevant clinical data and OS times were downloaded from their published studies^13,15–17^. The hazard ratios were calculated using the multivariate Cox proportional hazards model. All statistical significance testing in this study was two-sided. To control for multiple hypothesis testing, we applied the Benjamini-Hochberg method to correct p values and the false discovery rates (q-values) were calculated. Results were considered statistically significant at p-value or FDR q-value < 0.05.

### Data availability

All sequencing data generated during this study will be deposited in the Gene Expression Omnibus (GEO, https://www.ncbi.nlm.nih.gov/geo/). The data can be accessed under the accession number GSExxxxxx (the submission process is currently ongoing, and the accession number is going to be updated once it is complete).

## Acknowledgements

This study was supported in part by the IRG starts-up research funds provided to L.W. by U.T. MD Anderson Cancer Center (MDACC), the DOD grants: CA150334 and CA162445 to J.A.A., and DOD grants: CA160433 and CA170906 to S.S., the generous support from the Caporella, Dallas, Sultan, Park, Smith, Frazier, Oaks, Vanstekelenberg, Planjery, McNeil, Hyland, and Cantu families, as well as from the Schecter Private Foundation, Rivercreek Foundation, Kevin Fund, Myer Fund, Stupid Strong Foundation, Dio Fund, Milrod Fund, and the MDACC multidisciplinary grant programs. This study was also supported by SMF Core grant CA016672 (SMF). We thank E. J. Thompson, D. P. Pollock from the SMF Core for their excellent technical assistance. We thank all the patients who participated in this study.

## Author Contributions

L.W. and J.A. conceived and jointly supervised the study. S.S., K.H., M.P.P., M.Z., G.T., N.S., A.A.F.A., B.D.B., and M.B.M. contributed to sample collection and processing, and collection of patient clinical information. A.J.L., J.S.E., S.R.C. contributed to pathology review. L.W.supervised the bioinformatics data analysis, data integration and interpretation; R.W., contributed to sequencing data processing, quality check, integrative analyses, and generation of figures and tables for the manuscript. G.H., S.Z., Y.W., S.Z. assisted with data processing and analysis. L.W., J.A., R.W., A.J.L., P.A.F., S.H., G.A.C., and G.P. wrote and revised the manuscript.

## Competing Interest Statement

All authors declare no conflicts of interest.

**Extended Data Figures** (n=9, see the PDF file Extended-Data-Figures).

**Supplementary Tables** (n=5, see the PDF file Supplementary Tables_S1-4, and the Excel files Supplementary Tables_S5).

## Notes

**Conflicts of interest:** The authors have no potential conflicts of interest to disclose.

## REFERENCES

1. Bray, F., et al. Global cancer statistics 2018: GLOBOCAN estimates of incidence and mortality worldwide for 36 cancers in 185 countries. CA Cancer J Clin 68, 394–424 (2018).

2. Ikoma, N., et al. Preoperative chemoradiation therapy induces primary-tumor complete response more frequently than chemotherapy alone in gastric cancer: analyses of the National Cancer Database 2006-2014 using propensity score matching. Gastric Cancer 21, 1004–1013 (2018).

3. Mizrak Kaya, D., et al. Risk of peritoneal metastases in patients who had negative peritoneal staging and received therapy for localized gastric adenocarcinoma. J Surg Oncol 117, 678–684 (2018).

4. Shiozaki, H., et al. Prognosis of gastric adenocarcinoma patients with various burdens of peritoneal metastases. J Surg Oncol 113, 29–35 (2016).

5. Chen, C., et al. Efficacy and safety of immune checkpoint inhibitors in advanced gastric or gastroesophageal junction cancer: a systematic review and meta-analysis. Oncoimmunology 8, e1581547 (2019).

6. Taieb, J., et al. Evolution of checkpoint inhibitors for the treatment of metastatic gastric cancers: Current status and future perspectives. Cancer Treat Rev 66, 104–113 (2018).

7. Bartley, A.N., et al. HER2 Testing and Clinical Decision Making in Gastroesophageal Adenocarcinoma: Guideline From the College of American Pathologists, American Society for Clinical Pathology, and the American Society of Clinical Oncology. J Clin Oncol 35, 446–464 (2017).

8. Cancer Genome Atlas Research, N. Comprehensive molecular characterization of gastric adenocarcinoma. Nature 513, 202–209 (2014).

9. Wang, R., et al. Multiplex profiling of peritoneal metastases from gastric adenocarcinoma identified novel targets and molecular subtypes that predict treatment response. Gut (2019).

10. Tirosh, I., et al. Dissecting the multicellular ecosystem of metastatic melanoma by single-cell RNA-seq. Science 352, 189–196 (2016).

11. Jerby-Arnon, L., et al. A Cancer Cell Program Promotes T Cell Exclusion and Resistance to Checkpoint Blockade. Cell 175, 984–997 e924 (2018).

12. Puram, S.V., et al. Single-Cell Transcriptomic Analysis of Primary and Metastatic Tumor Ecosystems in Head and Neck Cancer. Cell 171, 1611–1624 e1624 (2017).

13. Patel, A.P., et al. Single-cell RNA-seq highlights intratumoral heterogeneity in primary glioblastoma. Science 344, 1396–1401 (2014).

14. canSAR (an integrated knowledge-base that provides drug-discovery useful predictions): https://cansarblack.icr.ac.uk.

15. Kim, H.K., et al. A gene expression signature of acquired chemoresistance to cisplatin and fluorouracil combination chemotherapy in gastric cancer patients. PLoS One 6, e16694 (2011).

16. Ooi, C.H., et al. Oncogenic pathway combinations predict clinical prognosis in gastric cancer. PLoS Genet 5, e1000676 (2009).

17. Cristescu, R., et al. Molecular analysis of gastric cancer identifies subtypes associated with distinct clinical outcomes. Nat Med 21, 449–456 (2015).

18. Mizrak Kaya, D., et al. Advanced gastric adenocarcinoma: optimizing therapy options. Expert Rev Clin Pharmacol 10, 263–271 (2017).

19. Dagogo-Jack, I. & Shaw, A.T. Tumour heterogeneity and resistance to cancer therapies. Nat Rev Clin Oncol 15, 81–94 (2018).

20. Hudler, P. Challenges of deciphering gastric cancer heterogeneity. World J Gastroenterol 21, 10510–10527 (2015).

21. Gullo, I., Carneiro, F., Oliveira, C. & Almeida, G.M. Heterogeneity in Gastric Cancer: From Pure Morphology to Molecular Classifications. Pathobiology 85, 50–63 (2018).

22. Oh, S.C., et al. Clinical and genomic landscape of gastric cancer with a mesenchymal phenotype. Nat Commun 9, 1777 (2018).

23. Merlo, L.M., Pepper, J.W., Reid, B.J. & Maley, C.C. Cancer as an evolutionary and ecological process. Nat Rev Cancer 6, 924–935 (2006).

24. Michor, F. & Polyak, K. The origins and implications of intratumor heterogeneity. Cancer Prev Res (Phila) 3, 1361–1364 (2010).

25. Amin, M.B., et al. AJCC cancer staging manual. 8th ed. New York: Springer (2017).

26. Amin, M.B., et al. The Eighth Edition AJCC Cancer Staging Manual: Continuing to build a bridge from a population-based to a more “personalized” approach to cancer staging. CA Cancer J Clin 67, 93–99 (2017).

27. Butler, A., Hoffman, P., Smibert, P., Papalexi, E. & Satija, R. Integrating single-cell transcriptomic data across different conditions, technologies, and species. Nat Biotechnol 36, 411–420 (2018).

28. Savas, P., et al. Single-cell profiling of breast cancer T cells reveals a tissue-resident memory subset associated with improved prognosis. Nat Med 24, 986–993 (2018).

29. Lambrechts, D., et al. Phenotype molding of stromal cells in the lung tumor microenvironment. Nat Med 24, 1277–1289 (2018).

30. Sade-Feldman, M., et al. Defining T Cell States Associated with Response to Checkpoint Immunotherapy in Melanoma. Cell 175, 998–1013 e1020 (2018).

31. Hanzelmann, S., Castelo, R. & Guinney, J. GSVA: gene set variation analysis for microarray and RNA-seq data. BMC Bioinformatics 14, 7 (2013).

32. Liu, J., et al. An Integrated TCGA Pan-Cancer Clinical Data Resource to Drive High-Quality Survival Outcome Analytics. Cell 173, 400–416 e411 (2018).

