## Extended Data Figures S1-9, and Supplementary Tables S1-4 for "Cell-of-Origin Analysis of Metastatic Gastric Cancer Uncovers the Origin of Inherent Intratumor Heterogeneity and a Fundamental Prognostic Signature"

**Extended Data Figure 1**

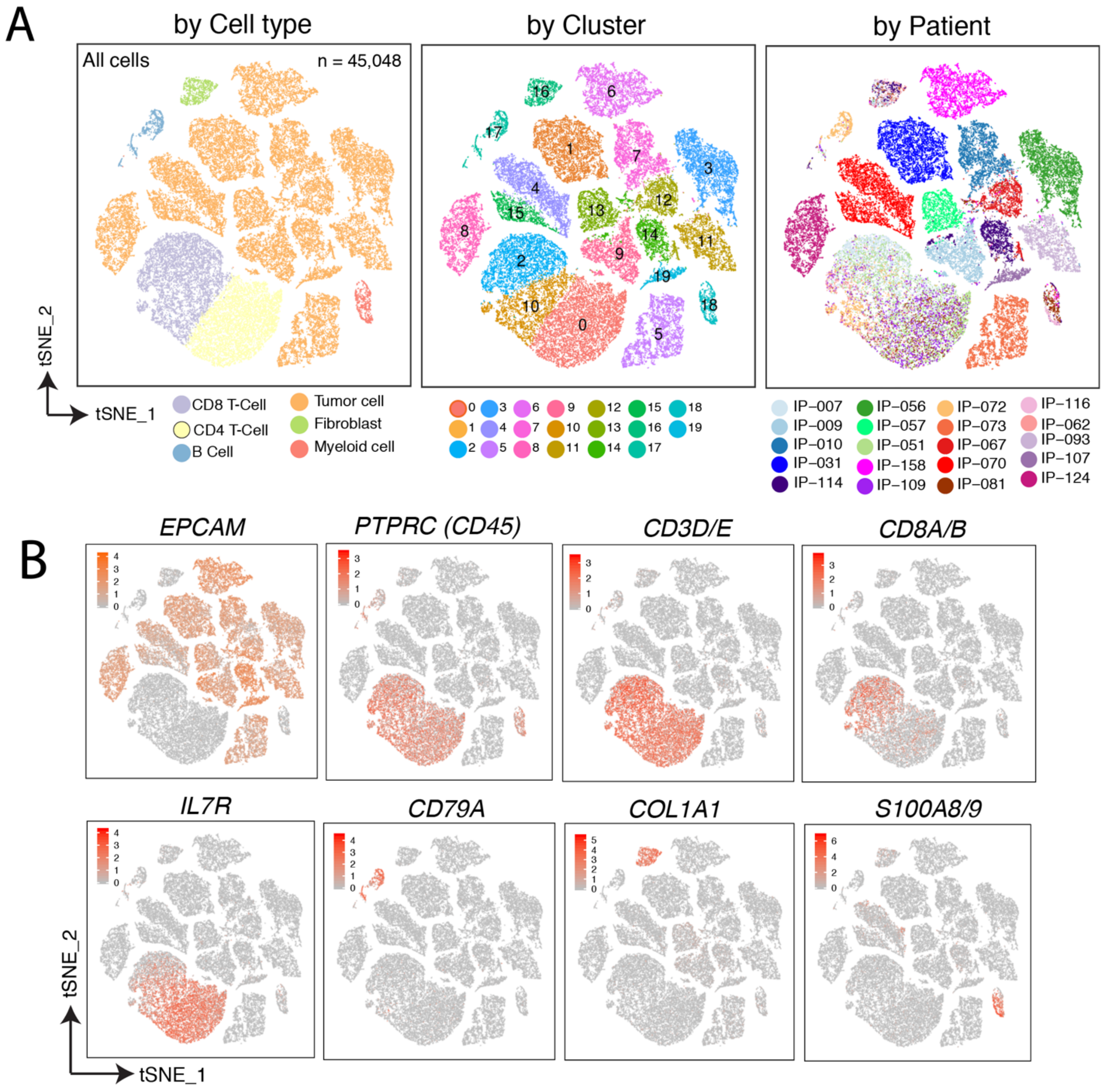


**A single cell transcriptome map of gastric peritoneal metastases**. Panel A shows an overview of the 45,048 single cells that passed quality control in this study. Each dot of the tSNE (t-distributed stochastic neighbor embedding) plot represents a single cell. Cells are color coded for (left to right): the associated cell type, tSNE cell cluster number, and the corresponding patient origin. Panel B shows the expression of canonical marker genes for the cell types defined above in the Panel A (left).

**Extended Data Figure 2**

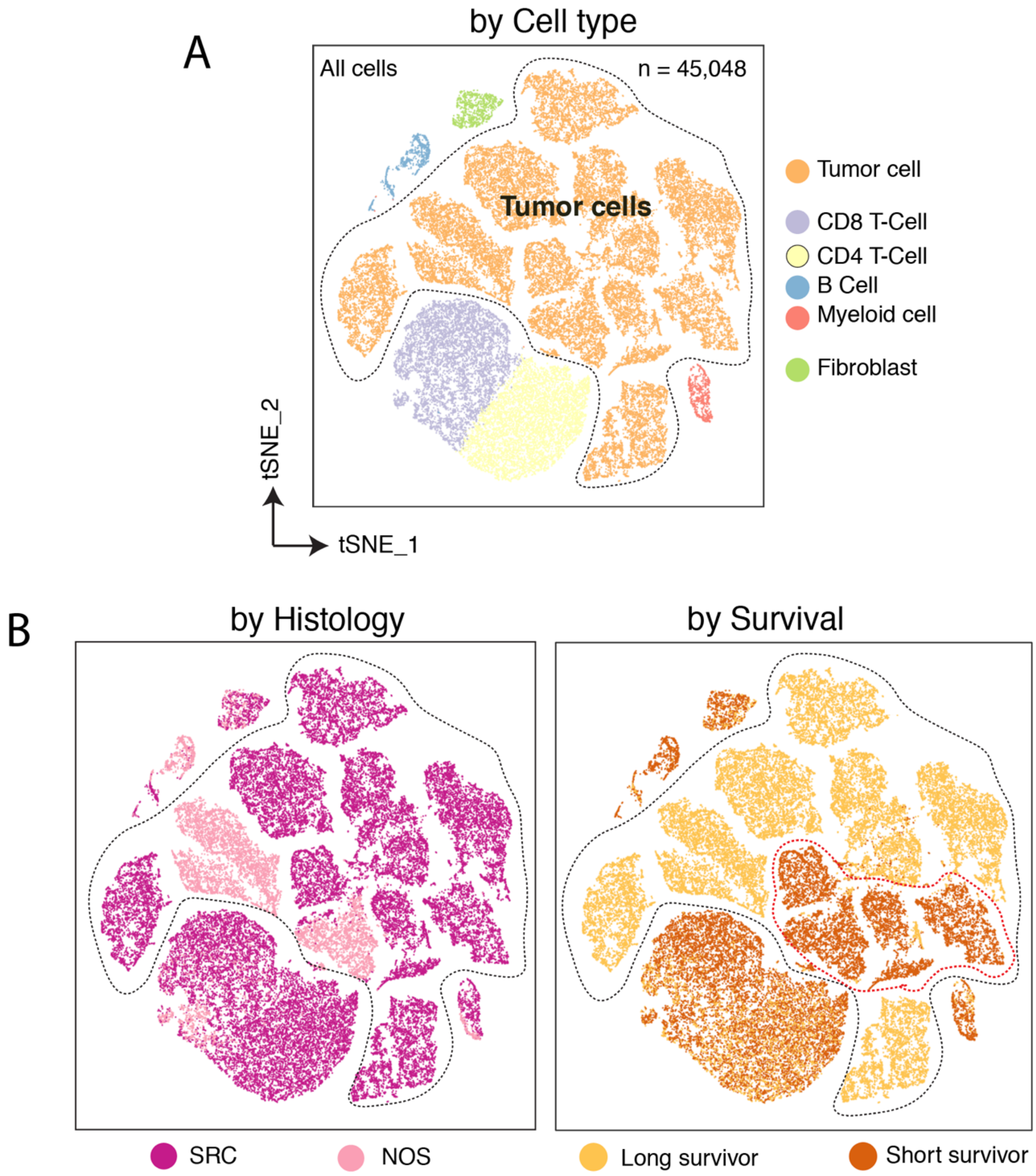


**Single cell clusters and correlation with histology types and clinical variables**. Panel A shows all cell clusters identified by unsupervised clustering. Tumor cells are highlighted by the black dashed irregular shape. Cells are color coded for the associated cell type (A), histology classification (B, left), and patient survival (B, right). SRC, Signet-ring cell carcinoma; NOS, not otherwise specified. The PC tumor cells from short survivors were clustered closely on the tSNE plot (B, right, highlighted by the red dashed irregular shape).

**Extended Data Figure 3**

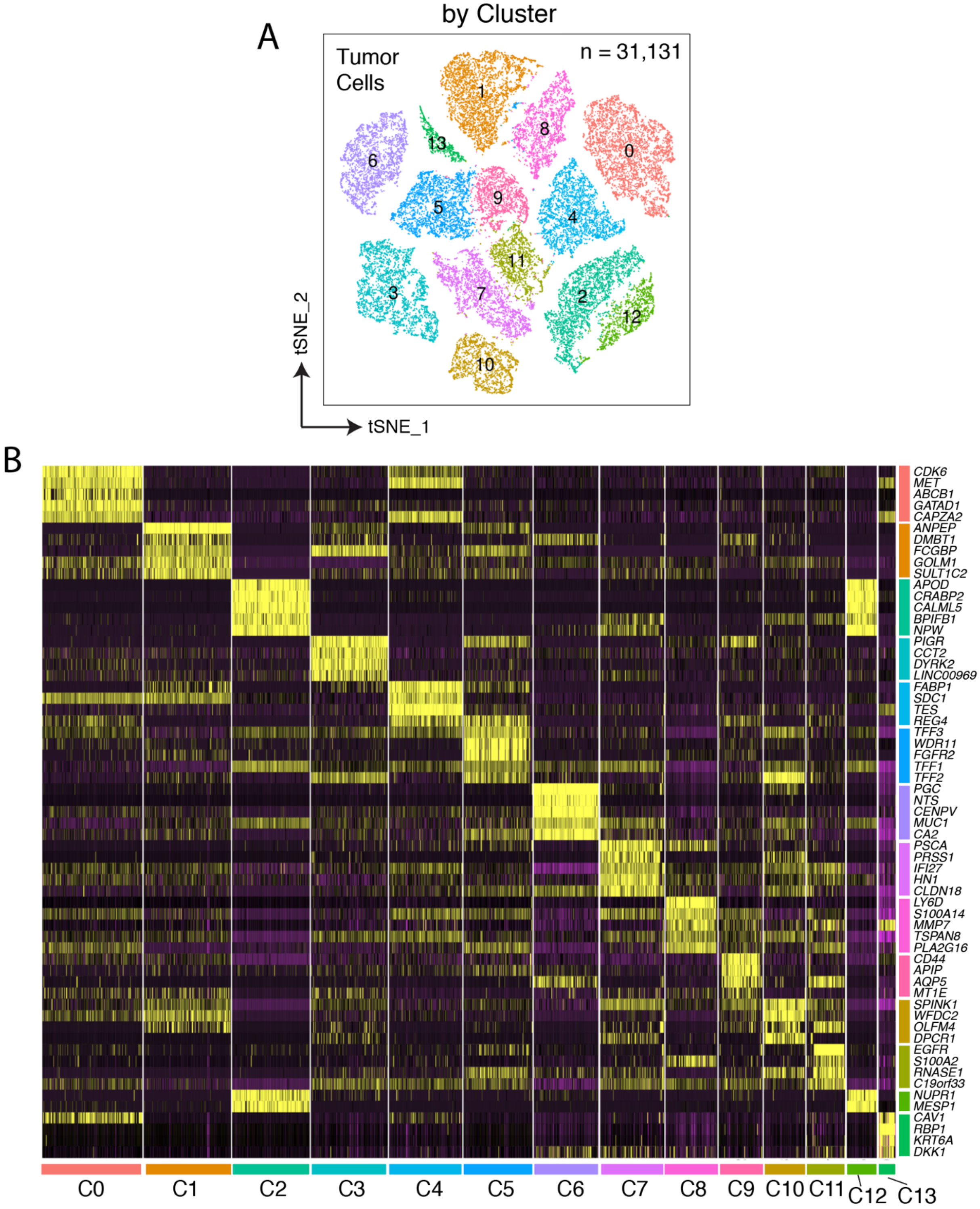


**Tumor cell heterogeneity at single cell level**. Panel A shows the tSNE overview of the 31,131 tumor cells that were passed QC and included for subsequent analysis. Cells formed 14 distinct clusters. The heatmap in Panel B displaying scaled expression values of discriminative genes (top 5 variable genes) per tumor cell cluster as defined in the panel A.

**Extended Data Figure 4**

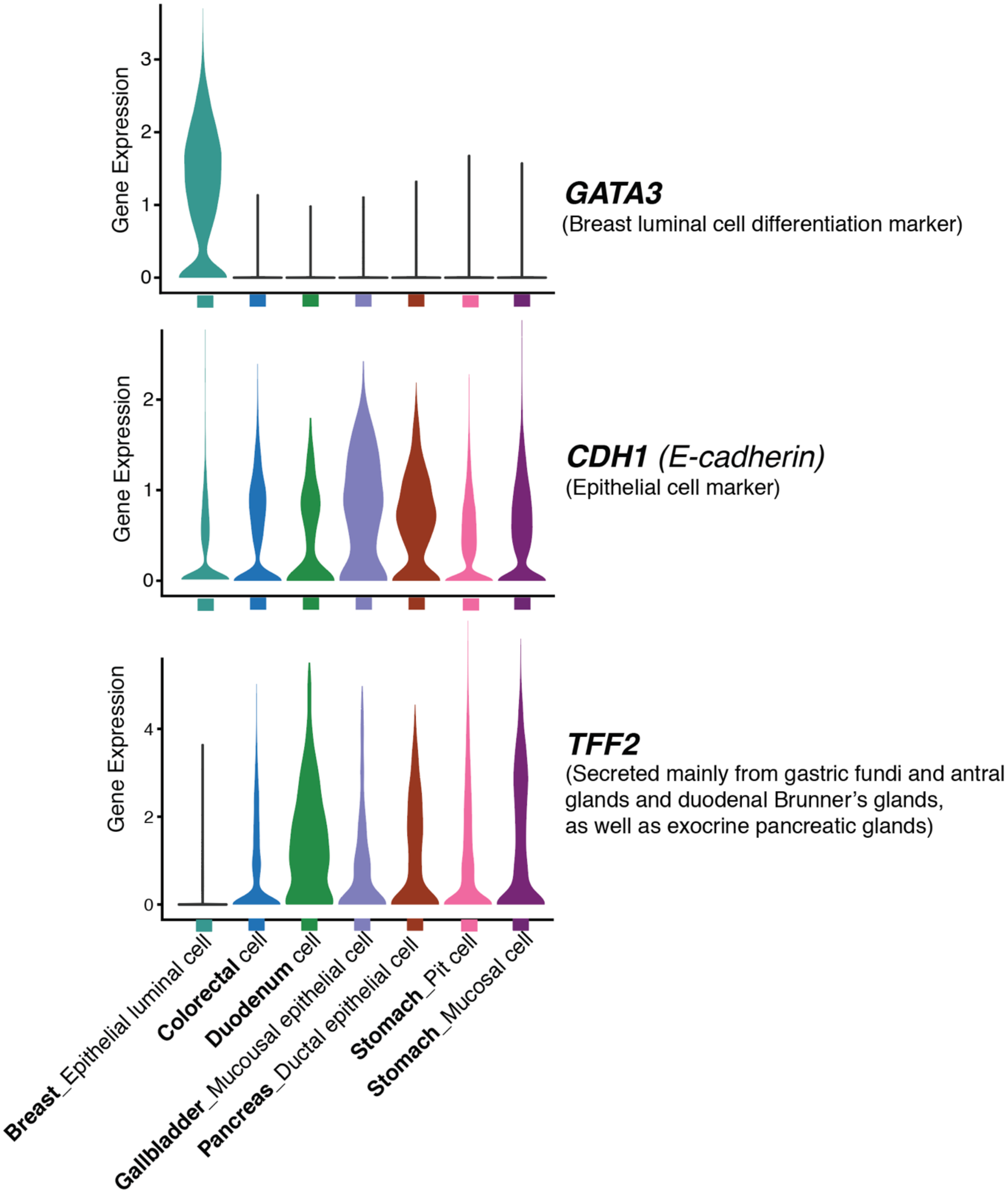


**Marker gene expression across tumor cells with different cell of origins**. The violin plots of 3 representative marker genes are shown. GATA3 is a differentiation marker for breast luminal epithelial cells; CDH1 (E-cadherin) is an epithelial cell marker; TFF2 is secreted mainly from the gastric glands, duodenal Brunner's glands, and exocrine pancreatic glands.

**Extended Data Figure 5**

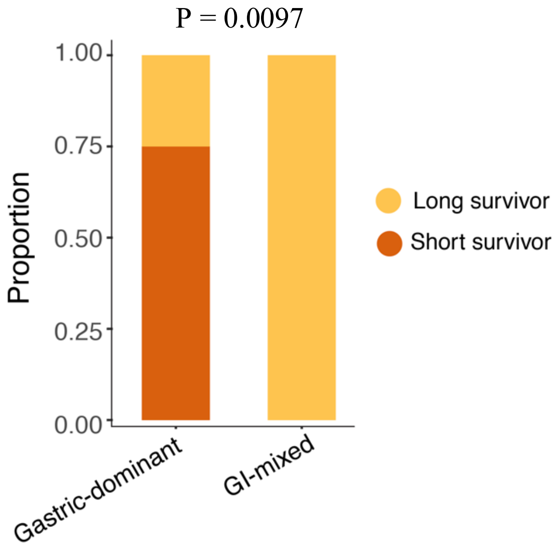


**Cell-of-origin based tumor classification was associated with patient survival**. The proportion of long and short-term survivors in cell-or-origin defined groups, the gastric dominant and the GI-mixed. P-value was calculated by two-tailed Fisher's Exact test.

**Extended Data Figure 6**

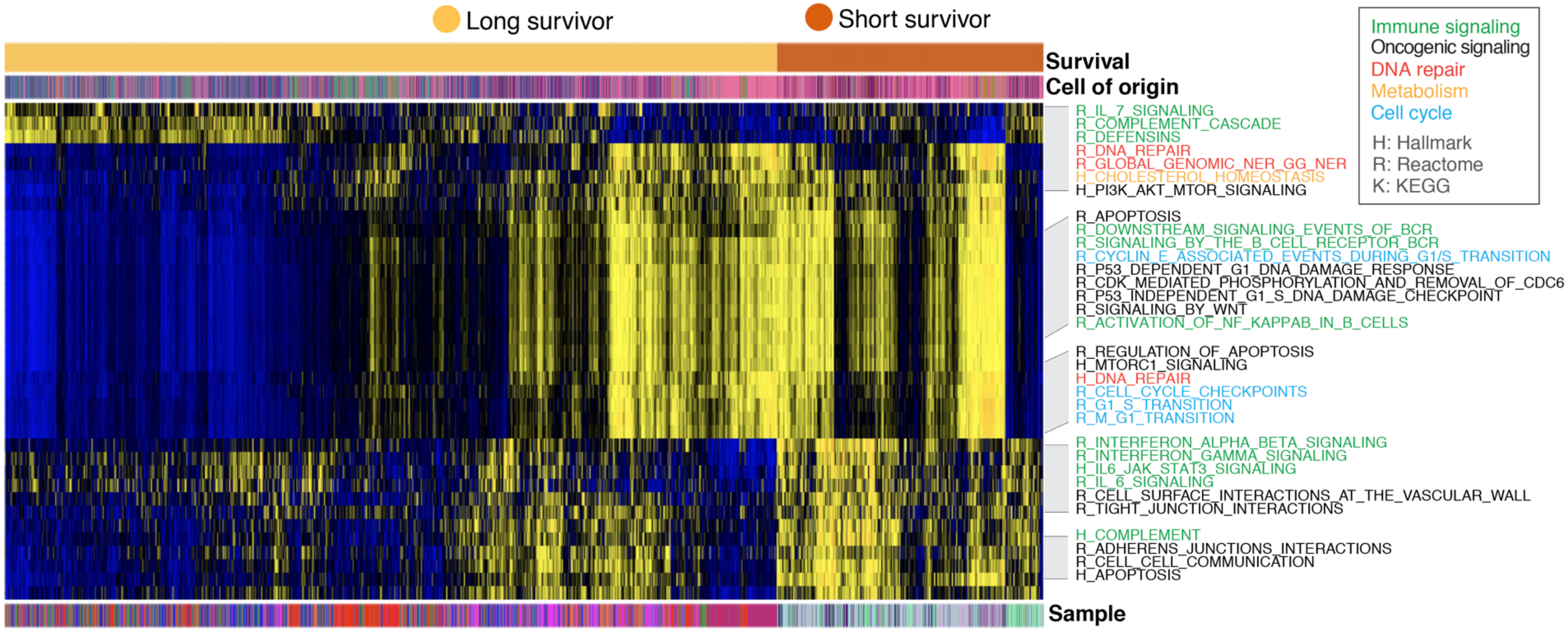


**Single cell signaling heterogeneity and correlation with patient survival**.
The pathways displayed in the Fig. 4A (differentially expressed across tumor cell origins) were included in the analysis and their pathway activity scores were compared between the short and long-term survivors and only those showed significant association with patient survivor are shown. The annotation track on the top labels tumor cells from long survivors (light brown) and short survivors (dark brown), respectively. Each row represents a single cell. The sample IDs were annotated at the bottom track using the same color code as shown

**Extended Data Figure 7**

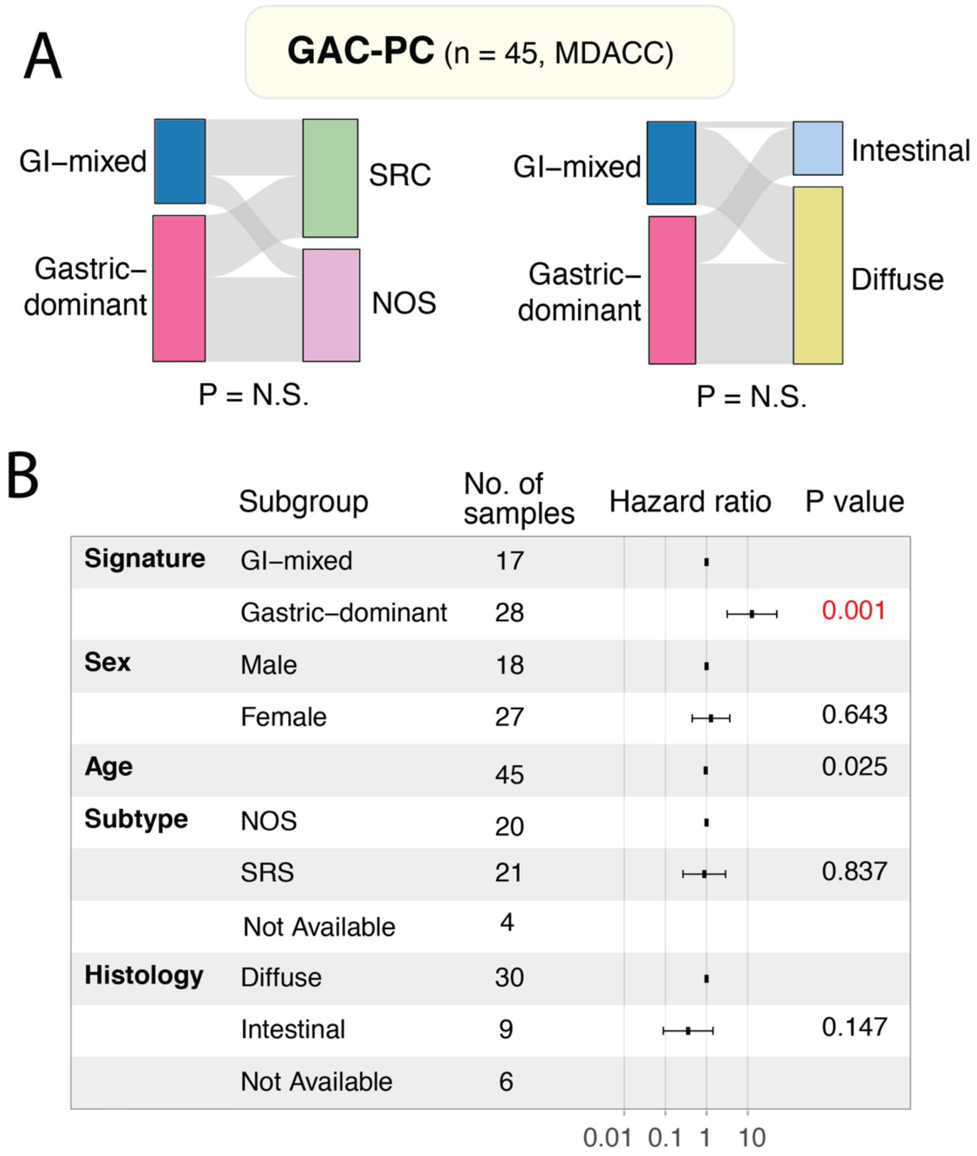


**The correlation of cell-of-origin defined subtypes with histology classifications and patient survival in an independent MDACC cohort of gastric peritoneal metastasis** (GAC-PC, n=45). The Alluvial plot in the Panel A shows the relationship between cell-of-origin defined subtypes and the histology based classifications: the presence of Signet-ring cell carcinoma (SRC) and the Lauren's classification of GAC. The left strip of each plot shows the cell-of-origin defined subtypes. The right strip shows the composition of histological subtypes. NOS, Not Otherwise Specified. N.S., not statistically significant. Panel B shows the multivariate analysis of the 12-gene signature in this patient cohort. The Cox proportional hazards regression model was used to calculate the hazard ration and P values.

**Extended Data Figure 8**

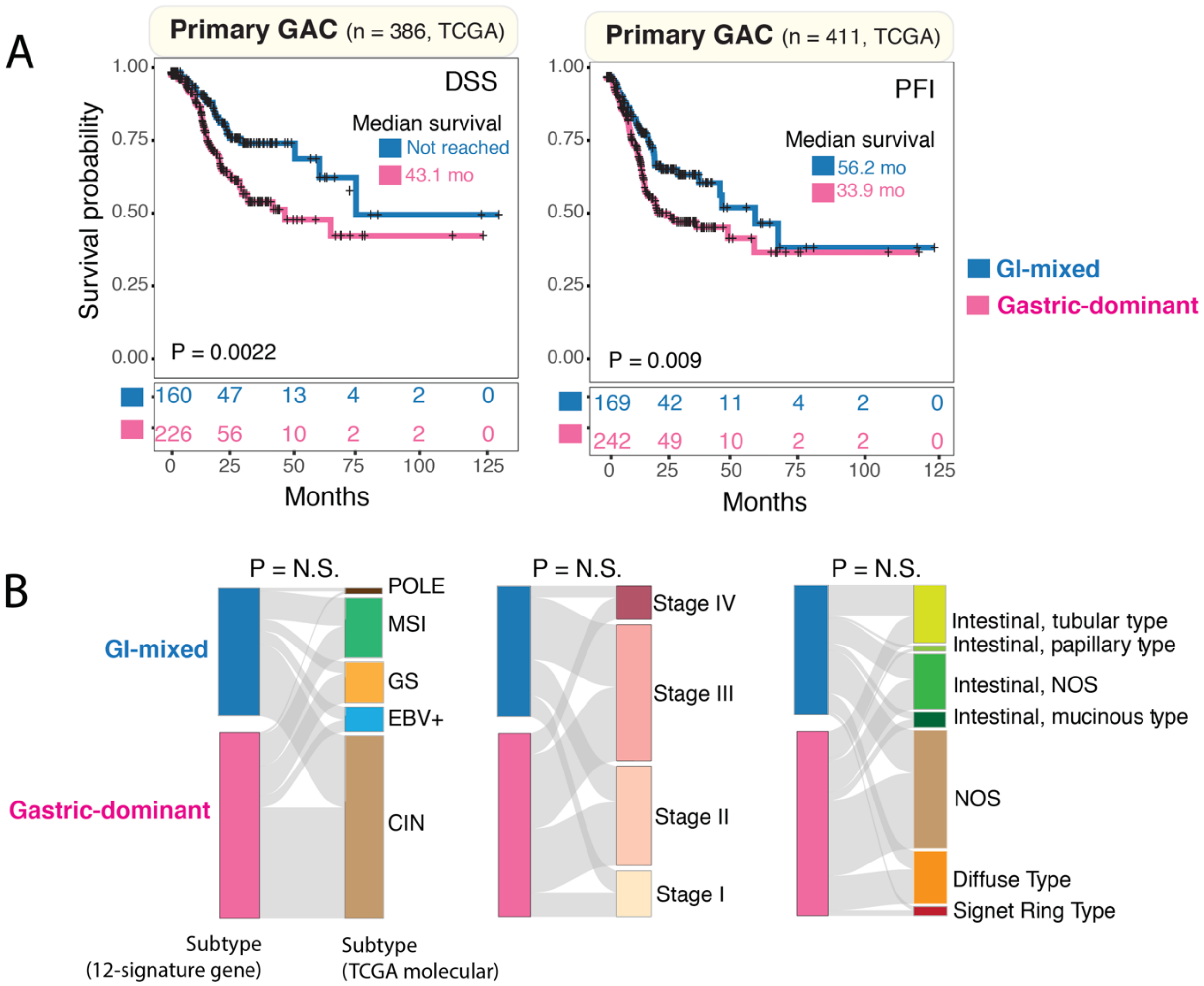


**The 12-gene signature and its prognostic significance in TCGA primary gastric cancer cohort and correlation with molecular subtypes and clinical variables**. The Kaplan–Meier curves in Panel A demonstrated the prognostic significance of the 12-gene signature in TCGA primary gastric cancer cohort (n=411). DSS, disease-specific survival (25 patients whose information were not available were excluded); PFI, progression free interval. The Alluvial plots in the Panel B show the relationship between cell-of-origin defined subtypes (left strip) and the molecular subtypes defined by TCGA multi-platform analysis (plot on the left, the right strip), tumor stage (plot in the middle, the right strip) and histology based classifications (plot on the right, the right strip). NOS, Not Otherwise Specified. N.S., not statistically significant.

**Extended Data Figure 9**

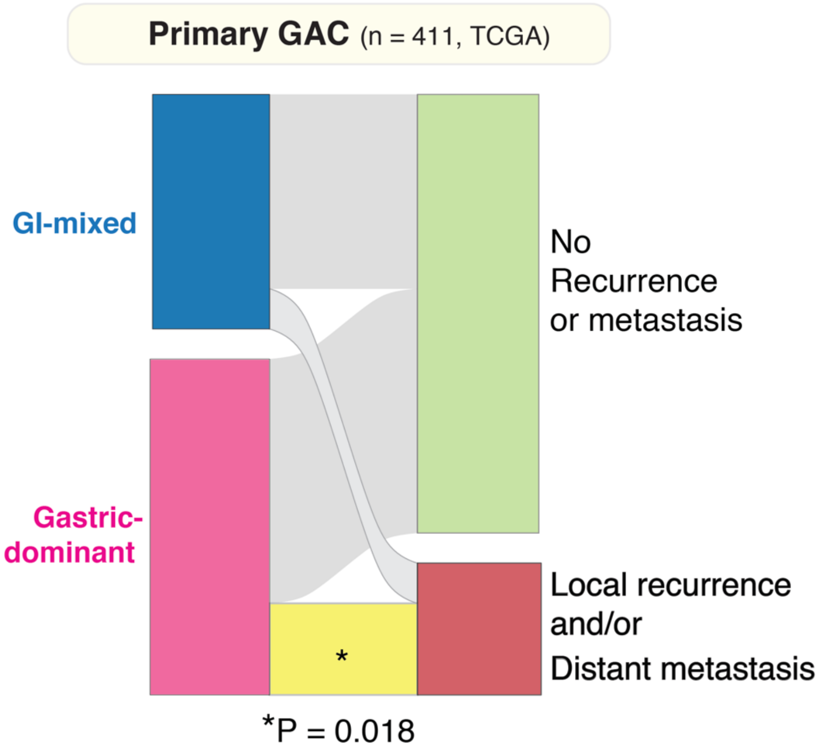


**The 12-gene signature and its prognostic significance in local disease recurrence and/or distal metastasis in the TCGA primary gastric cancer cohort** (n=411). The Alluvial plot shows the relationship between cell-of-origin defined subtypes (left strip) and the presence of local recurrence and/or distal metastasis (right strip). The yellow band highlights the significant enrichment of the local recurrence and/or distal metastasis events in tumors with the gastric-dominant subtype.

### SUPPLEMENTARY TABLES

**Table S1.**

**The clinical and histopathological characteristics of 20 GAC-PC patients.**

| Patient ID | Survival group | Status | Survival (from peritoneal metastasis) (month) | Race | Gender | Tumor Grade | The Lauren's Classification | Signet-ring cell carcinoma | Intestinal Metaplasia | Her2 Positivity | H. pylori | Primary Site |
|---|---|---|---|---|---|---|---|---|---|---|---|---|
| IP-007 | Short | Dead | 4.5 | Arabic | Male | G3 | Diffuse | Yes | N/A | No | Yes | Distal |
| IP-009 | Short | Dead | 5.0 | White | Male | G3 | Diffuse | No | N/A | No | No | Proximal |
| IP-010 | Long | alive | 14.7 | White | Female | G3 | Diffuse | Yes | Yes | No | No | Distal |
| IP-031 | Long | Dead | 19.9 | Asian | Male | G3 | Diffuse | Yes | N/A | No | N/A | Proximal |
| IP-051 | Short | Dead | 4.8 | Black | Female | G3 | Diffuse | Yes | N/A | No | No | Distal |
| IP-056 | Long | Dead | 21.8 | White | Female | G3 | Diffuse | Yes | N/A | No | N/A | Distal |
| IP-057 | Short | Dead | 5.2 | White | Male | G3 | Diffuse | Yes | No | N/A | No | Proximal |
| IP-062 | Long | alive | 44.9 | White | Female | G3 | Diffuse | Yes | N/A | No | No | Distal |
| IP-067 | Long | Dead | 18.1 | White | Female | G3 | Diffuse | Yes | N/A | No | N/A | Distal |
| IP-070 | Long | Dead | 17.3 | Hispanic | Female | G3 | Diffuse | No | N/A | No | No | Distal |
| IP-072 | Short | Dead | 1.6 | White | Male | G3 | Diffuse | No | N/A | No | No | Proximal |
| IP-073 | Long | Dead | 22.1 | Hispanic | Male | G3 | Diffuse | Yes | No | No | No | Proximal |
| IP-081 | Short | Dead | 3.3 | Black | Female | G3 | Diffuse | Yes | N/A | No | No | Distal |
| IP-093 | Short | Dead | 5.6 | Black | Female | G3 | Diffuse | Yes | N/A | N/A | No | Distal |
| IP-107 | Short | Dead | 2.8 | Hispanic | Female | G3 | Diffuse | Yes | N/A | No | No | Distal |
| IP-109 | Long | Dead | 16.4 | White | Female | G3 | Diffuse | Yes | N/A | No | N/A | Distal |
| IP-114 | Short | Dead | 0.4 | Black | Male | G3 | Diffuse | Yes | N/A | N/A | N/A | remnant gastric |
| IP-116 | Short | Dead | 0.4 | White | Male | G3 | Diffuse | No | N/A | No | No | Proximal |
| IP-124 | Long | Dead | 33.8 | White | Female | G3 | Diffuse | Yes | N/A | No | No | Distal |
| IP-158 | Long | alive | 24.9 | Hispanic | Female | G3 | Diffuse | Yes | Yes | No | N/A | Distal |

Note: N/A, not available

*Note: Long survivor, patients survived > 1 year after the diagnosis of peritoneal metastasis; Short survivor, patients died with 6 months after the diagnosis of peritoneal metastasis (see more details in Supplementary Methods).*

### Table S2.

### A summary matrix showing the number of cells with defined cell of origins in each sample.

| Cell of Origin (QC-passed tumor cells) | IP-093 | IP-124 | IP-009 | IP-114 | IP-081 | IP-057 | IP-109 | IP-007 | IP-073 | IP-056 | IP-158 | IP-010 | IP-031 | IP-067 | IP-070 | IP-051 | IP-062 | IP-072 | IP-107 | IP-116 |
|---|---|---|---|---|---|---|---|---|---|---|---|---|---|---|---|---|---|---|---|---|
| *Stomach_Mucosal cell* | 405 | 73 | 1098 | 69 | 39 | 274 | 17 | 16 | 774 | 335 | 1168 | 1305 | 154 | 81 | 0 | 0 | 1 | 0 | 0 | 0 |
| *Stomach_Pit cell* | 1382 | 2230 | 1055 | 855 | 25 | 927 | 66 | 28 | 1361 | 1956 | 527 | 547 | 801 | 573 | 1 | 2 | 0 | 0 | 0 | 5 |
| *Stomach_Chief cell* | 0 | 74 | 0 | 0 | 0 | 0 | 0 | 1 | 0 | 51 | 0 | 2 | 0 | 0 | 0 | 0 | 0 | 0 | 0 | 0 |
| *Pancreas_Ductal cell* | 8 | 3 | 38 | 155 | 11 | 284 | 0 | 9 | 89 | 74 | 4 | 1 | 292 | 50 | 0 | 2 | 6 | 0 | 1 | 10 |
| *Gallbladder_Mucous epithelial cell* | 2 | 0 | 8 | 9 | 10 | 26 | 1 | 0 | 0 | 2 | 65 | 2 | 120 | 32 | 0 | 1 | 1 | 0 | 0 | 6 |
| *Duodenum_Enterocyte progenitor* | 0 | 0 | 0 | 0 | 0 | 0 | 0 | 1 | 44 | 52 | 0 | 4 | 0 | 0 | 0 | 0 | 0 | 0 | 0 | 0 |
| *Duodenum_Goblet cell* | 0 | 30 | 0 | 0 | 0 | 31 | 6 | 1 | 23 | 148 | 0 | 13 | 1 | 11 | 0 | 1 | 0 | 0 | 0 | 0 |
| *TransverseColon_Goblet cell* | 0 | 0 | 0 | 4 | 2 | 0 | 0 | 1 | 171 | 247 | 251 | 469 | 1023 | 488 | 2 | 0 | 0 | 0 | 0 | 0 |
| *TransverseColon_Enterocyte* | 0 | 0 | 1 | 0 | 0 | 0 | 0 | 2 | 47 | 11 | 416 | 80 | 166 | 1 | 0 | 0 | 0 | 0 | 0 | 0 |
| *TransverseColon_Enterocyte progenitor* | 0 | 0 | 0 | 0 | 0 | 0 | 0 | 3 | 27 | 86 | 20 | 49 | 93 | 40 | 0 | 0 | 0 | 0 | 0 | 0 |
| *SigmoidColon_Enterocyte progenitor* | 0 | 0 | 0 | 0 | 0 | 0 | 0 | 3 | 162 | 148 | 116 | 27 | 262 | 145 | 0 | 0 | 0 | 0 | 0 | 1 |
| *Rectum_Enterocyte* | 0 | 0 | 0 | 0 | 0 | 0 | 0 | 1 | 108 | 15 | 104 | 59 | 62 | 6 | 0 | 0 | 0 | 0 | 0 | 0 |
| *Rectum_Inflammed epithelial cell* | 1 | 0 | 0 | 0 | 0 | 2 | 0 | 0 | 47 | 30 | 3 | 6 | 224 | 46 | 0 | 0 | 0 | 0 | 0 | 0 |
| *Breast_Epithelium_luminal cell* | 0 | 0 | 0 | 0 | 0 | 0 | 0 | 0 | 0 | 0 | 0 | 0 | 0 | 0 | 3631 | 0 | 0 | 0 | 0 | 0 |
| *Other* | 98 | 11 | 50 | 143 | 9 | 20 | 14 | 5 | 12 | 142 | 77 | 18 | 537 | 61 | 382 | 3 | 2 | 1 | 650 | 21 |
| Total tumor cells | 1896 | 2421 | 2250 | 1235 | 96 | 1564 | 104 | 71 | 2865 | 3297 | 2751 | 2582 | 3735 | 1534 | 4016 | 9 | 10 | 1 | 651 | 43 |
| Tumor tumor cells with defined cell of origin | 1798 | 2410 | 2200 | 1092 | 87 | 1544 | 90 | 66 | 2853 | 3155 | 2674 | 2564 | 3198 | 1473 | 3634 | 6 | 8 | 0 | 1 | 22 |
| Included in subsequent analsysis | Yes | Yes | Yes | Yes | Yes | Yes | Yes | Yes | Yes | Yes | Yes | Yes | Yes | Yes | Yes | No | No | No | No | No |

*Note: The sample IDs are shown on the top with each column represent an individual sample. The defined cell types (Supplementary Methods) are shown on the left (row). Other, other unclassified cells or super rare cell types. Five samples with less than 50 tumor cells with defined cell of origins in each sample were excluded from subsequent analysis (gray colored on the right).*

**Table S3.**

**A summary matrix showing the number of cells with defined cell cycle stages in each sample.**

| Sample ID | G0/G1 | G2M | S | Total cells |
|---|---|---|---|---|
| IP-093 | 1055 | 283 | 558 | 1,896 |
| IP-124 | 657 | 690 | 1074 | 2,421 |
| IP-009 | 973 | 749 | 528 | 2,250 |
| IP-114 | 511 | 403 | 321 | 1,235 |
| IP-081 | 75 | 7 | 14 | 96 |
| IP-057 | 1095 | 206 | 263 | 1,564 |
| IP-109 | 54 | 22 | 28 | 104 |
| IP-007 | 36 | 16 | 19 | 71 |
| IP-073 | 1615 | 606 | 644 | 2,865 |
| IP-056 | 1139 | 909 | 1249 | 3,297 |
| IP-158 | 1904 | 299 | 548 | 2,751 |
| IP-010 | 1174 | 800 | 608 | 2,582 |
| IP-031 | 1992 | 945 | 798 | 3,735 |
| IP-067 | 512 | 454 | 568 | 1,534 |
| IP-070 | 2163 | 1057 | 796 | 4,016 |

*Note: the cell cycle stage was computationally assigned based on the expression profiles of cell cycle related genes (Supplementary Methods).*

### Table S4

**The genes amplified by 17q copy number gain and significantly associated with patient survival.**

| Symbol | logFC | Expression in Cells from Short Survivor (mean) | Expression in Cells from Long Survivor (mean) | P-value | FDR q-value |
|---|---|---|---|---|---|
| NOTCH1 | 1.36 | 1.00 | 0.89 | 0 | 0 |
| H3F3B | 1.12 | 1.00 | 0.98 | 0 | 0 |
| SUMO2 | 0.81 | 0.98 | 0.83 | 0 | 0 |
| MRPL12 | 0.74 | 0.93 | 0.76 | 0 | 0 |
| ATP5H | 0.73 | 0.98 | 0.83 | 0 | 0 |
| ANAPC11 | 0.69 | 0.99 | 0.81 | 0 | 0 |
| STRA13 | 0.69 | 0.94 | 0.72 | 0 | 0 |
| LGALS3BP | 0.68 | 0.99 | 0.90 | 0 | 0 |
| DCXR | 0.67 | 0.94 | 0.66 | 0 | 0 |
| MRPS7 | 0.61 | 0.88 | 0.52 | 0 | 0 |
| ARHGDIA | 0.58 | 0.97 | 0.74 | 0 | 0 |
| NME1 | 0.57 | 0.89 | 0.72 | 0 | 0 |
| RPL38 | 0.56 | 1.00 | 0.99 | 0 | 0 |
| SYNGR2 | 0.54 | 0.97 | 0.84 | 0 | 0 |
| P4HB | 0.50 | 1.00 | 0.93 | 0 | 0 |
| UBALD2 | 0.50 | 0.81 | 0.45 | 0 | 0 |
| PHB | 0.47 | 0.93 | 0.75 | 0 | 0 |
| SAP30BP | 0.46 | 0.84 | 0.38 | 0 | 0 |
| C17orf89 | 0.46 | 0.91 | 0.67 | 0 | 0 |
| ACTG1 | 0.46 | 1.00 | 0.99 | 0 | 0 |
| SLC9A3R1 | 0.43 | 0.80 | 0.51 | 0 | 0 |
| SLC16A3 | 0.42 | 0.84 | 0.47 | 0 | 0 |
| NARF | 0.40 | 0.82 | 0.43 | 0 | 0 |
| PSMC5 | 0.39 | 0.89 | 0.61 | 0 | 0 |
| SEPTIN9 | 0.38 | 0.89 | 0.58 | 0 | 0 |
| SNF8 | 0.38 | 0.85 | 0.58 | 0 | 0 |
| EIF1 | 0.38 | 1.00 | 0.97 | 0 | 0 |
| GPS1 | 0.37 | 0.82 | 0.49 | 0 | 0 |
| SIRT7 | 0.34 | 0.82 | 0.53 | 0 | 0 |
| DDX5 | 0.31 | 0.99 | 0.90 | 0 | 0 |
| SOCS3 | 0.31 | 0.64 | 0.24 | 0 | 0 |
| WBP2 | 0.30 | 0.70 | 0.31 | 0 | 0 |
| EIF4A3 | 0.30 | 0.71 | 0.46 | 0 | 0 |
| CYTH1 | 0.29 | 0.58 | 0.19 | 0 | 0 |
| GRB2 | 0.28 | 0.74 | 0.39 | 0 | 0 |
| EXOC7 | 0.27 | 0.72 | 0.35 | 0 | 0 |
| PRPSAP1 | 0.27 | 0.64 | 0.27 | 0 | 0 |
| TSEN54 | 0.26 | 0.68 | 0.33 | 0 | 0 |
| RNF213 | 0.26 | 0.81 | 0.48 | 0 | 0 |
| TMC6 | 0.26 | 0.70 | 0.34 | 0 | 0 |
| SRP68 | 0.26 | 0.61 | 0.25 | 0 | 0 |
| UNC13D | 0.25 | 0.52 | 0.16 | 0 | 0 |
| ATP5G1 | 0.53 | 0.93 | 0.81 | 5E-297 | 1E-292 |
| MRPL27 | 0.26 | 0.85 | 0.65 | 9E-261 | 2E-256 |
| PSMB3 | 0.28 | 0.93 | 0.81 | 1E-231 | 3E-227 |
| RPL27 | 0.37 | 1.00 | 0.95 | 1E-200 | 3E-196 |
| RPL19 | 0.36 | 1.00 | 0.99 | 2E-154 | 4E-150 |
| SLC25A39 | 0.25 | 0.83 | 0.73 | 6E-118 | 1E-113 |
| VMP1 | -0.34 | 0.93 | 0.86 | 2E-16 | 4E-12 |

*Note: Only the genes located to 17q copy number amplification (Fig. 3B) were included in analysis and their expression profiles were compared between the samples from long and short-term survivors and the ones significantly associated with patient survivor are listed. FC, fold change; FDR q-value are adjusted p values for multiple hypothesis testing.*
